# Resistance to the KRAS^G12D^ Inhibitor MRTX1133 is Associated with Increased Sensitivity to BET Inhibition

**DOI:** 10.1101/2025.05.10.653074

**Authors:** Daniel R. Principe, Jeffrey H. Becker, Anastasia E. Metropulos, Alejandra M. Marinelarena, Thao D. Pham, Alexandre F. Aissa, Hidayatullah G. Munshi

## Abstract

As many as 90% of human pancreatic ductal adenocarcinoma (PDAC) tumors harbor gain-of-function mutations in the *KRAS* oncogene. Recently, inhibitors of the most common KRAS mutation, KRAS^G12D^, have entered the clinical arena. However, early evidence suggests that as monotherapy, KRAS^G12D^ inhibitors such as MRTX1133 at best provide brief periods of disease stabilization. Hence, there is a growing interest in understanding the mechanisms through which tumors acquire resistance to KRAS inhibition. In the present study, we generated *in vitro* models of MRTX1133 resistance and subjected parental and drug-resistant cell lines to RNA sequencing. This suggested that MRTX1133-resistant tumor cells undergo a global shift toward histone acetylation. Inhibition of the histone acetyltransferase EP300 reversed the drug-resistant phenotype *in vitro*, which subsequent RNA sequencing experiments determined was associated with the suppression of pro-survival FOSL1 signaling. Accordingly, *siFOSL1* reversed the MRTX1133-resistant phenotype with similar effects on pro-survival signaling. Given the lack of clinically useful EP300 or FOSL1 inhibitors, we next explored whether inhibitors of the acetylation scanning BET proteins would be similarly effective. The addition of BET inhibitors re-sensitized several resistant cell lines to MRTX1133 and impaired FOSL1-mediated survival signaling *in vitro*. In murine models of MRTX1133-resistant PDAC, BET inhibition cooperated with MRTX1133 to markedly extend overall survival. As BET inhibitors are currently under clinical testing, the combination of MRTX1133 and BET inhibitors warrants further investigation, particularly in tumors that have developed resistance to KRAS inhibition.

**SIGNIFICANCE:** Here, we demonstrate that BET inhibition is effective in PDAC with acquired resistance to KRAS inhibitors. As BET inhibitors are under clinical testing, the combination of KRAS and BET inhibitors warrants consideration in PDAC patients.

**Graphical Abstract:** 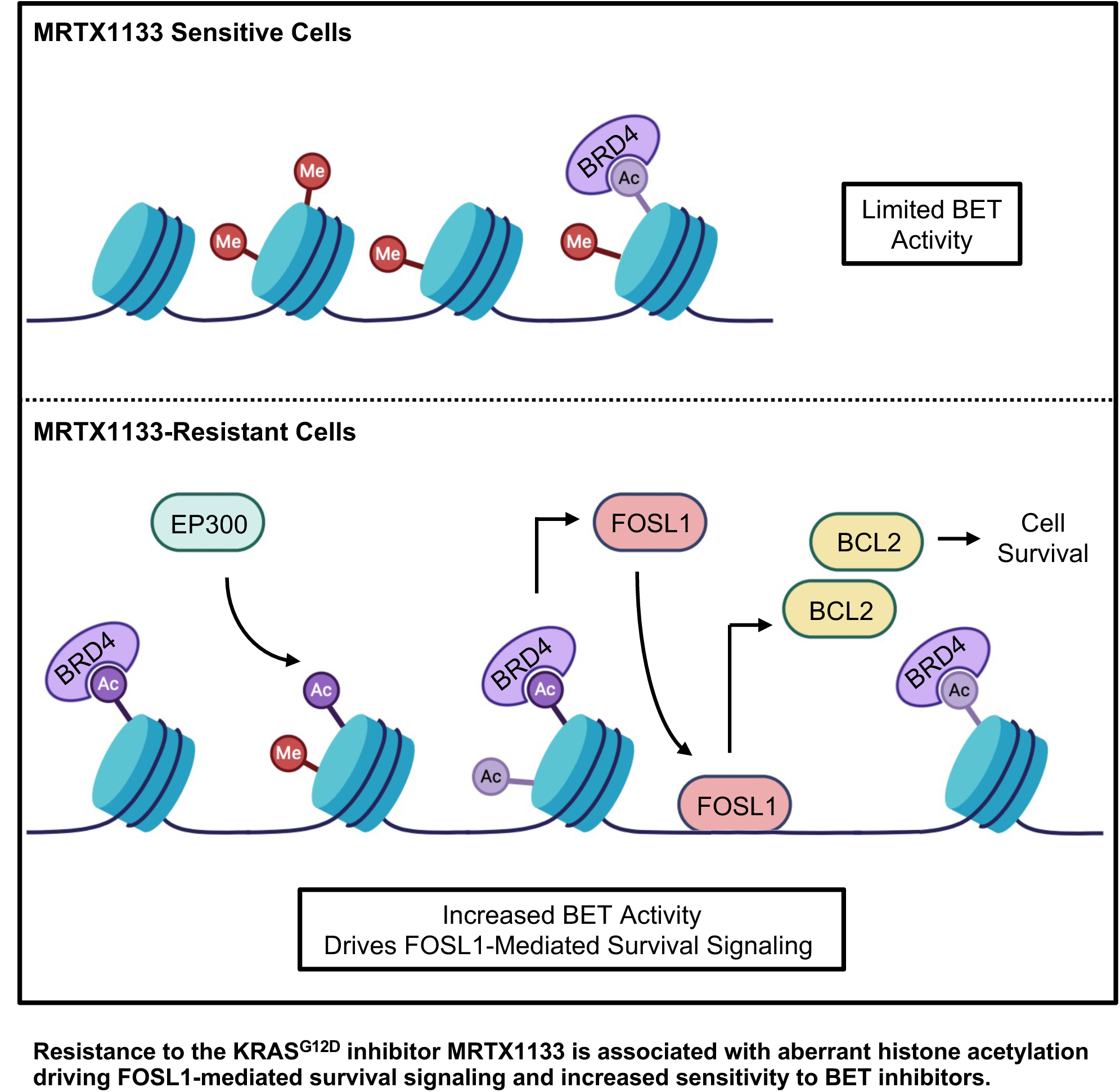

## INTRODUCTION

Pancreatic ductal adenocarcinoma (PDAC) is projected to become the second leading cause of cancer-related death by 2030 (1). Despite significant advances in chemo-, immuno-, and targeted therapy, nearly all PDAC tumors will develop resistance to the current standard of care treatments, and overall survival remains a dismal 13% (2,3). Gain-of-function *KRAS* mutations are present in over 90% of human PDAC tumors, and oncogenic KRAS has central roles in tumor initiation and maintenance (4). Accordingly, mutant KRAS has long been suggested as an attractive therapeutic target in PDAC tumors. The KRAS^G12C^ inhibitor Sotorasib was the first selective KRAS inhibitor approved by the Federal Drug Administration (FDA) based on data from a phase 3 clinical trial of KRAS^G12C^-mutated non-small cell lung cancer (NSCLC) patients (5). As *KRAS^G12C^* mutations are present in 1-2% of human PDAC tumors, selective KRAS^G12C^ inhibitors have also been tested in PDAC patients (6,7). However, single-agent treatment with KRAS^G12C^ inhibitors at best led to brief periods of disease stabilization in PDAC patients (7,8).

Recently, selective inhibitors of the most common KRAS mutation (KRAS^G12D^) have also entered the clinical arena. While the KRAS^G12D^ inhibitor MRTX1133 has shown promise in preclinical studies, like clinical data regarding KRAS^G12C^ inhibitors, MRTX1133 treatment in mice provided only transient periods of disease stabilization followed by progression (9,10). As MRTX1133 is now in the early phase of clinical testing, there is a growing interest in identifying novel drug combinations to improve therapeutic efficacy, and several potential strategies are now emerging (11). For example, in xenograft tumor models, the effects of MRTX1133 are enhanced by the addition of either EGFR or AKT inhibition (12). Similarly, in immunocompetent mice, the combination of MRTX1133 and immune checkpoint inhibitors (ICIs) enhanced therapeutic responses (13,14). Though these strategies have shown promise, in several studies, tumors eventually show signs of progression, indicative of acquired resistance.

Though several studies have explored resistance to KRAS^G12C^ inhibitors (5,15,16), the cellular mechanisms underlying resistance to KRAS^G12D^ inhibitors are still emerging (17). Given the promise of MRTX1133 and other KRAS^G12D^ inhibitors in PDAC patients, there is a pressing need for additional therapeutic strategies in tumors that have developed resistance to MRTX1133. To address this, we first generated two *in vitro* models of MRTX1133 resistance and subjected both parental and drug-resistant cell lines to RNA sequencing. Though the transcriptomes of the two MRTX1133-resistant cell lines differed, both had significant alteration to several gene sets associated with epigenetic processes suggestive of increased histone acetylation.

After confirming a hyperacetylated phenotype of the MRTX1133-resistant tumor cells, we next incubated drug-resistant cells with a pharmacologic inhibitor of the histone acetyltransferase (HAT) EP300. Interestingly, EP300-inhibition reversed the MRTX1133-resistant phenotype and impaired FOSL1-mediated survival signaling. Consistent with these observations, siRNA against *FOSL1* reversed the MRTX1133-resistant phenotype and inhibited pro-survival signaling. Finally, given the lack of clinically useful EP300 or FOSL1 inhibitors, we hypothesized that inhibitors of the acetylation scanning BET proteins may be similarly effective. Though BET inhibitors did not enhance the efficacy of MRTX1133 in naïve cell lines, the addition of BET inhibitors sensitized several highly resistant cell lines to MRTX1133 and impaired FOSL1-mediated survival signaling *in vitro*. In animal models, the addition of the BET inhibitor JQ1 re-sensitized resistant tumors to MRTX1133. This led to improved overall survival, downregulation of FOSL1, reduced proliferation, and increased cell death. As BET inhibitors are currently under clinical testing, the combination of MRTX1133 and BET inhibitors warrants further investigation, particularly in tumors that have developed resistance to KRAS inhibition.

## MATERIALS & METHODS

### Antibodies and Key Reagents

All antibodies were purchased from established commercial vendors and verified by the manufacturer for the specific species and applications for which they were used. MRTX1133 and JQ1 for *in vitro* use were purchased from SelleckChem (SelleckChem, Houston, TX), and for *in vivo* use from MedChemExpress (Monmouth, Junction, NJ) and MedKoo Biosciences, Inc. (Durham, NC). For a full list of antibodies used, please see **Table S1**.

### Cell Culture, siRNA, and Viability Assays

Human PDAC cell lines PANC1, ASPC1, and CD18 were purchased from the ATCC. Murine PDAC cell lines KPC-2138 (2138) and KPC-3213 (3213) were derived from the *LSL-Kras**^G12D/+^**/LSL-Trp53**^R172H/+^**/Pdx-1-Cre* (KPC) model in the C57BL/6 background as previously described (18). The FC1245 cell line was provided by Dr. David Tuveson (Cold Spring Harbor Laboratories, Cold Spring Harbor, NY), and was also generated from the KPC model in the C57BL/6 background (19). All cell lines were maintained in DMEM supplemented with 10% Fetal Bovine Serum (FBS), 100 U/mL Penicillin, and 100 µg/mL Streptomycin (ThermoFisher, Waltham, MA). To generate MRTX1133-resistant cell lines, each of the above cell lines were incubated in increasing concentrations of MRTX1133 until viable in ≥2µM, after which cells were maintained in 2µM MRTX1133 (20). All cell lines were used within 15 passages after revival from frozen stocks and tested for mycoplasma every six months via the LookOut Mycoplasma PCR Detection Kit (Sigma Aldrich, St. Louis, MO) and used if negative.

For transfections, silence-select pre-designed siRNAs against either human *FOSL1* or murine *Fosl1* were obtained from DharmaCon (DharmaCon, Lafayette, CO). Scramble siRNAs were used as controls (Santa Cruz, Dallas, TX). The siRNA transfections were carried out using RNAiMAX (Invitrogen, Waltham, MA) per the manufacturer’s instructions. For cell viability assays, 2,000-4,000 cells were seeded into each well of a 96-well plate in serum-free DMEM. After 16 hours, media/drug was added and cells cultured for 72 hours. At this time, we added 20 μL of a 5mg/ml 3-(4,5-Dimethyl-2-thiazolyl)-2,5-diphenyl-2H-tetrazolium bromide (MTT) solution to each well (1:100). After two hours, media was aspirated, crystals dissolved in DMSO, and 570 nm absorbance determined by plate reader. All experiments were performed in triplicate unless otherwise specified.

### Floating Collagen Gel Cultures

Floating collagen gel cultures were established as described previously (20). In brief, acidified rat tail Collagen I (Corning, Corning, NY) was neutralized with 0.34N NaOH and diluted to a final concentration of 1.2 mg/ml. Cells were trypsinized, and single-cell suspensions mixed with the diluted collagen mixture to obtain a final concentration of 1×10^5^ cells/mL. The collagen/cell mixture suspension was vortexed, and 1 mL seeded into each well of an Ultra-Low Attachment 6-well plates (Corning) as described (21,22). The mixture was allowed to polymerize at 37°C for one hour, after which 2mL of culture media was added and gels detached from the well using a fine pipette tip. Cells were imaged using the EVOS XL Core Imaging System (ThermoFisher), and the relative growth was quantified as described previously (20).

### Western Blot

Cell or tissue lysates were lysed in radioimmunoprecipitation assay buffer (RIPA) buffer (ThermoFisher) followed by sonication. Equal amounts of protein (15–50 μg) were mixed with loading dye, boiled for 8 min, separated on a 4-20% denaturing SDS–polyacrylamide gel electrophoresis (PAGE) gel and transferred to a Polyvinylidene difluoride (PVDF) membrane. The membrane was blocked in 5% milk/TBS/0.1% Tween for one hour and incubated with antibodies against Ac-H3-K9, BCL2 (human), pERK, ERK, FOSL1 (mouse), pMEK, MEK (Cell Signaling, Danvers, MA), Ac-H3-K56 (Active Motif, Carlsbad, CA), KRAS (Novus Bio, Saint Charles, MO), Ac-H4, Pan Histone (Milipore, Temecula, CA), FOSL1 (human), Ac-H3-K27 (abcam, Cambridge, MA), BCL2 (mouse) (ThermoFisher), or GAPDH (Santa Cruz Biotech, Santa Cruz, CA). The membrane was washed with tris-buffered saline (TBS)-0.1% Tween and then incubated with horseradish peroxidase (HRP) conjugated secondary antibody (Cell Signaling) at room temperature for one hour and rewashed. Protein bands were visualized by an enhanced chemiluminescence method (ThermoFisher) and resolved digitally per the manufacturer’s specifications.

### RNA sequencing and Gene Set Enrichment Analysis

PANC1, PANC1K, 2138, or 2138K cells were seeded into floating collagen gel cultures and treated as described. RNA was extracted using the RNeasy Plus Mini Kit (Qiagen, Hilden, Germany) per the manufacturer’s instructions. Quality control, sequencing, and data processing were performed by ActiveMotif. The raw count files were processed using DESeq2 (23) with default parameters to identify differentially expressed genes between cells treated with the molecules of interest and control cells. For Gene Set Enrichment Analysis (GSEA) (24), shrunken log2 fold change results from DESeq2 were ranked according to log2 fold change and analyzed using the ClusterProfiler package (25) against GMT files for C5 (Gene Ontology) Biological Process from the MSigDB collections, utilizing human or mouse datasets (Human MSigDB v2023.2.Hs, Mouse MSigDB v2023.2.Mm) (26,27). Only enriched terms with adjusted p-values (p.adjust) < 0.05 were reported. Heatmaps of individual genes were selected from the core enrichment GSEA terms, and the top, most significantly altered genes with the lowest p-values were plotted using the pheatmap package (Version 1.0.12) in R (28). Data are in the process of being deposited under accession number GSE294216.

### qPCR

Quantitative gene expression was performed with gene-specific TaqMan probes, TaqMan Universal PCR Master Mix, and the 7500 Fast Real-time PCR System from Applied Biosystems (Foster City, CA). These data were quantified with the comparative cycle threshold (CT) method for relative gene expression.

### Orthotopic Tumor Models

To establish 1245K-derived tumors, 10,000 cells were collected in 10 µL of DMEM and mixed with 10 µL of growth factor reduced, LDEV-free basement membrane matrix/Matrigel (Corning). The cell suspension was kept on ice until immediately prior to use. The entire 20 µL suspension was injected into the pancreas of background-matched C57BL/6 mice, and mice were allowed to develop 200-250 mm^3^ tumors. At this time, mice were randomized into one of four treatment groups. Mice were treated with daily, intraperitoneal injections of DMSO (vehicle control), 30 mg/kg MRTX1133 (10,12,14), 50 mg/kg JQ1 (29,30), or a combination of MRTX1133 and JQ1 (N = 10 mice/group). 2138K-derived tumors were established similarly, using a total of 7,500 cells. Once animals developed large 800-1,000 mm^3^ tumors, they were enrolled into one of four treatment groups as described above. All mice were euthanized either upon ulceration of the skin or when showing clear signs of health decline, e.g., weight loss, ascites, or lethargy.

### Subcutaneous Tumor Models

To establish subcutaneous 1245, 1245K, 2318, and 2138K-derived tumors, 25,000 cells were collected as above, and subcutaneously injected into the flanks of 6-8-week-old C57BL/6 mice. Mice were euthanized when tumor volume exceeded 1200-1500 mm^3^, ulcerated, or when mice showed clear signs of health decline, e.g., weight loss, ascites, or lethargy.

### Histology, Immunohistochemistry, and Immunofluorescence

Mice were euthanized and the pancreas, colon, small bowel, liver, and spleen were subjected to pathologic examination. Tissues were fixed in 10% formalin, paraffin-embedded, and sections at 4 mm interval were cut from each tissue and stained with hematoxylin and eosin (H&E), or via immunohistochemistry (IHC) or immunofluorescence (IF). For IHC, slides were deparaffinized by xylenes and rehydrated by ethanol gradient, then heated in a pressure cooker using sodium citrate buffer (10mM sodium citrate, 0.05% Tween 20, pH 6.0). Tissues were blocked with 0.5% bovine serum albumin (BSA) in PBS for 30 minutes and incubated with primary antibodies against BCL2 (abcam), pERK, FOSL1, or Cleaved Caspase 3 (Cell Signaling) at 1:100–1:1000 overnight at 4°C. Slides were developed using HRP conjugated secondary antibodies followed by 3,3′-Diaminobenzidine (DAB) substrate/buffer (DAKO).

For IF, slides were heated via pressure cooker in sodium citrate buffer and tissues blocked with 0.5% BSA in PBS for 1 hour at room temperature. Sections were exposed to primary antibodies against CK19 (University of Iowa Hybridoma Bank) or Ki67 (Cell Signaling) at 1:100-1:1000 overnight at 4°C. Slides were developed using AlexaFluor 488-or 647-conjugated secondary antibodies (1:500-1:1,000, ThermoFisher), mounted in DAPI-containing media (Santa Cruz Biotechnology), exposed to DAPI, Fluorescein isothiocyanate (FITC), and Cy5 filters.

### Microscopy

All brightfield images were acquired using a Leica DM2500 LED optical microscope with bright field camera attachment. Negative slides were used for white balance, and for all images no analog or digital gain was used. For fluorescent imaging, images were acquired using the EVOS M700 imaging system (ThermoFisher) microscope with fluorescent camera attachment.

### Tissue Slide Counts and Measurements

All counts were performed by a minimum of two blinded investigators and each value displayed includes the average of at least three high power fields per specimen as described in our previous publications (31–35). Score distributions were visualized via Minitab express software, showing the median value as a solid line and all individual values excluding any statistical outliers.

### Statistical analysis

Non-sequencing, non-survival data were analyzed by two-way ANOVA and fit to a general linear model in Minitab16, the validity of which was tested by adherence to the normality assumption and the fitted plot of the residuals. Results were arranged by the Tukey method and were considered significant at p < 0.05. Results are presented as individual value plots showing the median value as a solid line and all individual values excluding any statistical outliers unless otherwise noted. Survival data was analyzed by the Kaplan-Meier/log rank test method as described previously (36).

### Study approval

All experiments involving the use of mice were performed following protocols approved by the IACUC at Northwestern University.

### Data availability

All raw data generated in this study are available upon request from the corresponding authors.

## RESULTS

### MRTX1133 resistance is associated with the upregulation of several epigenetic processes

To identify potential mechanisms of resistance to KRAS^G12D^ inhibitors, we first generated two *in vitro* models of MRTX1133 resistance by incubating the human PDAC cell line PANC1 or the murine PDAC cell line KPC-2138 (2138) in increasing concentrations of MRTX1133. Once cells were viable in a minimum concentration of 2µM, cells were referred to as PANC1K and 2138K, respectively (**Figure 1A**). Once these cells were established, the drug-resistant phenotype was first confirmed by Western blotting, showing relative insensitivity to MRTX1133-induced MEK/ERK inactivation compared to parental cells (**Figure 1B,C and S1A**). We next serum-starved parental and MRTX1133-resistant cells overnight, after which cells were changed to full-serum media containing either a DMSO vehicle or increasing concentrations of MRTX1133. After another 72 hours, cell viability was evaluated by 3-(4,5-Dimethylthiazol-2-yl)-2,5-diphenyltetrazolium bromide (MTT) assay. This showed an expected decline in viability for both parental cell populations, yet no significant change in cell viability for either PANC1K or 2138K cells (**Figure 1D,E**). As PDAC cells are more sensitive to the effects of MRTX1133 when grown in 3D floating collagen gels (20), both parental and MRTX1133-resistant cells were next seeded at low density in 3D floating collagen cultures and incubated with either a DMSO vehicle or a fixed 0.5 µM dose of MRTX1133. Cell confluence was quantified over 6 days, and consistent with our previous observations (20), treatment with MRTX1133 led to pronounced growth suppression in parental PANC1 and 2138 cells, yet failed to significantly alter the growth kinetics of PANC1K and 2138K cells (**Figure 1F,G**).

**Figure 1.**
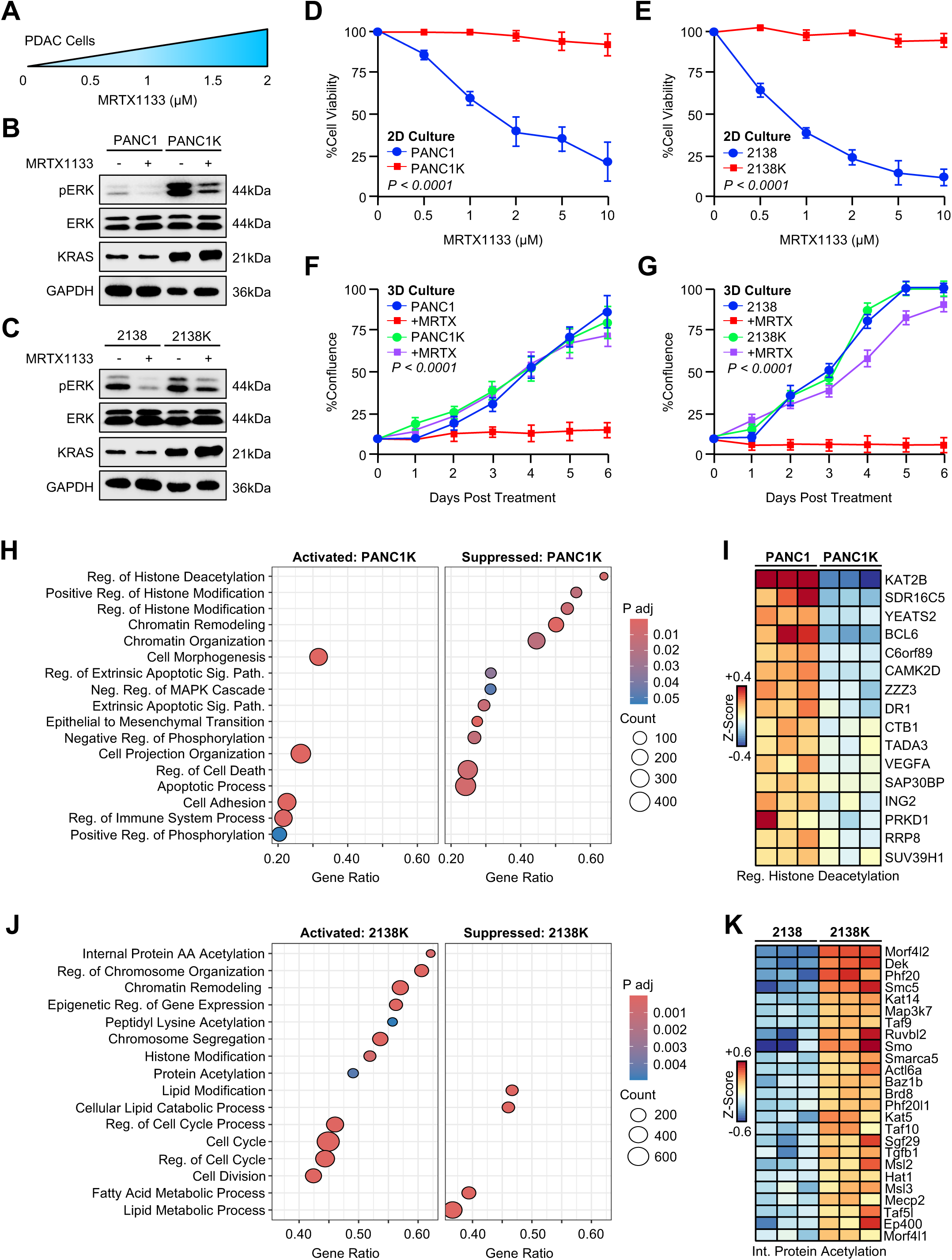
MRTX1133 resistance is associated with the upregulation of several epigenetic processes **(A)** PANC1 or 2138 tumor cells were incubated with increasing concentrations of MRTX1133 until viable in 2μM. After this point, cells were referred as PANC1K or 2138K for KRAS-inhibitor resistant cells. (**B,C**) Parental and KRAS-inhibitor resistant cells were serum-starved overnight, pre-treated with 2µM MRTX1133 for 30 minutes, after which they were changed to full serum media containing 2µM MRTX1133. After 24 hours, pERK was evaluated by western blot. **(D)** PANC1 or PANC1K cells were collected in serum-free media and seeded into 96-well plates (4,000/cells per well). Cells were changed to full-serum media containing either a DMSO vehicle or increasing concentrations of MRTX1133. After another 72 hours, cell viability was evaluated by 3-(4,5-Dimethylthiazol-2-yl)-2,5-diphenyltetrazolium bromide (MTT) assay. **(E)** 2138 or 2138K cells were seeded in serum-free media into 96-well plates as previously. The next morning, media was changed to full-serum media containing either a DMSO vehicle or increasing concentrations of MRTX1133 and cell viability evaluated by MTT assay after 72 hours. **(F)** PANC1 or PANC1K cells were seeded in 3D floating collagen cultures (10,000 cells/well) and incubated with either a DMSO vehicle or a fixed 0.5 µM dose of MRTX1133. Cell confluence was quantified, and growth shown over a period of 6 days. **(G)** 2138 or 2138K cells were seeded in 3D floating collagen gels (10,000 cells/well) and incubated with either a DMSO vehicle or a fixed 0.5 µM dose of MRTX1133. Cell confluence was quantified over 6 days. **(H,I)** PANC1 and PANC1K cells were seeded in 3D floating collagen cultures for 48 hours and subjected to RNA sequencing and gene set enrichment analysis (GSEA) performed. Significantly altered gene sets are shown using a Z-score cutoff of 2.5. Focused heatmap is shown for select, significantly altered genes in the regulation of histone deacetylation gene set. **(J,K)** 2138 and 2138K cells were seeded in 3D floating collagen and also subjected to RNA sequencing and GSEA. Focused heatmap is shown for select, significantly altered genes in the internal protein acetylation gene set.

After verifying the MRTX1133-resistant phenotype, PANC1 and PANC1K cells were again grown in 3D floating collagen cultures for 48 hours and subjected to RNA sequencing followed by gene set enrichment analysis (GSEA). This identified several cell processes differentially expressed between the two cell lines. Of note, PANC1K cells had significant downregulation of several genes involved in epigenetic cell processes, namely those involved in the histone deacetylation pathway. Consistent with the observed increase in ERK phosphorylation, PANC1K cells also demonstrated suppression of negative regulators of MAPK signaling, as well as downregulation in several genes involved in cell death/apoptotic signaling (**Figure 1H,I**). We repeated this experiment using 2138 and 2138K cells, with GSEA identifying a significant upregulation in several gene sets related to histone modification and chromatin organization. Additionally, 2138K cells had suppression of select lipid metabolic processes and enhanced expression of several genes associated with cell cycle progression and cell proliferation (**Figure 1J,K**).

### Inhibition of EP300-mediated histone acetylation reverses MRTX1133 resistance and reduces FOSL1 expression

Though the transcriptomes of PANC1K and 2138K cells had several differences, both had significant changes in epigenetic gene sets, namely those relating to histone acetylation. These data suggest that PANC1K cells have a relative loss of deacetylase activity, whereas 2138K cells have enhanced histone acetyltransferase activity. Both cases suggest a global shift toward histone acetylation in the MRTX1133-resistant cell lines. We, therefore, evaluated relative changes in histone acetylation by Western blot. This demonstrated a relative increase in histone acetylation (H3-K27, H3-K56, H4) in PANC1K and 2138K cells compared to their parental cell line **(Figure 2A,B)**. Given our group’s previous findings that the histone acetyltransferase EP300 functions as a central regulator of oncogenic KRAS signaling in PDAC (31), we next sought to determine whether inhibition of EP300-mediated histone acetylation would re-sensitize these resistant cell lines to MRTX1133. We therefore incubated PANC1K or 2138K cells with either a DMSO vehicle or 5µM of the EP300 inhibitor SGC-CBP30. After 72 hours, cells were seeded into 96-well plates, and in addition to DMSO or SGC-CBP30, media was supplemented with increasing concentrations of MRTX1133. After another 72 hours, cell viability was evaluated by MTT assay.

**Figure 2.**
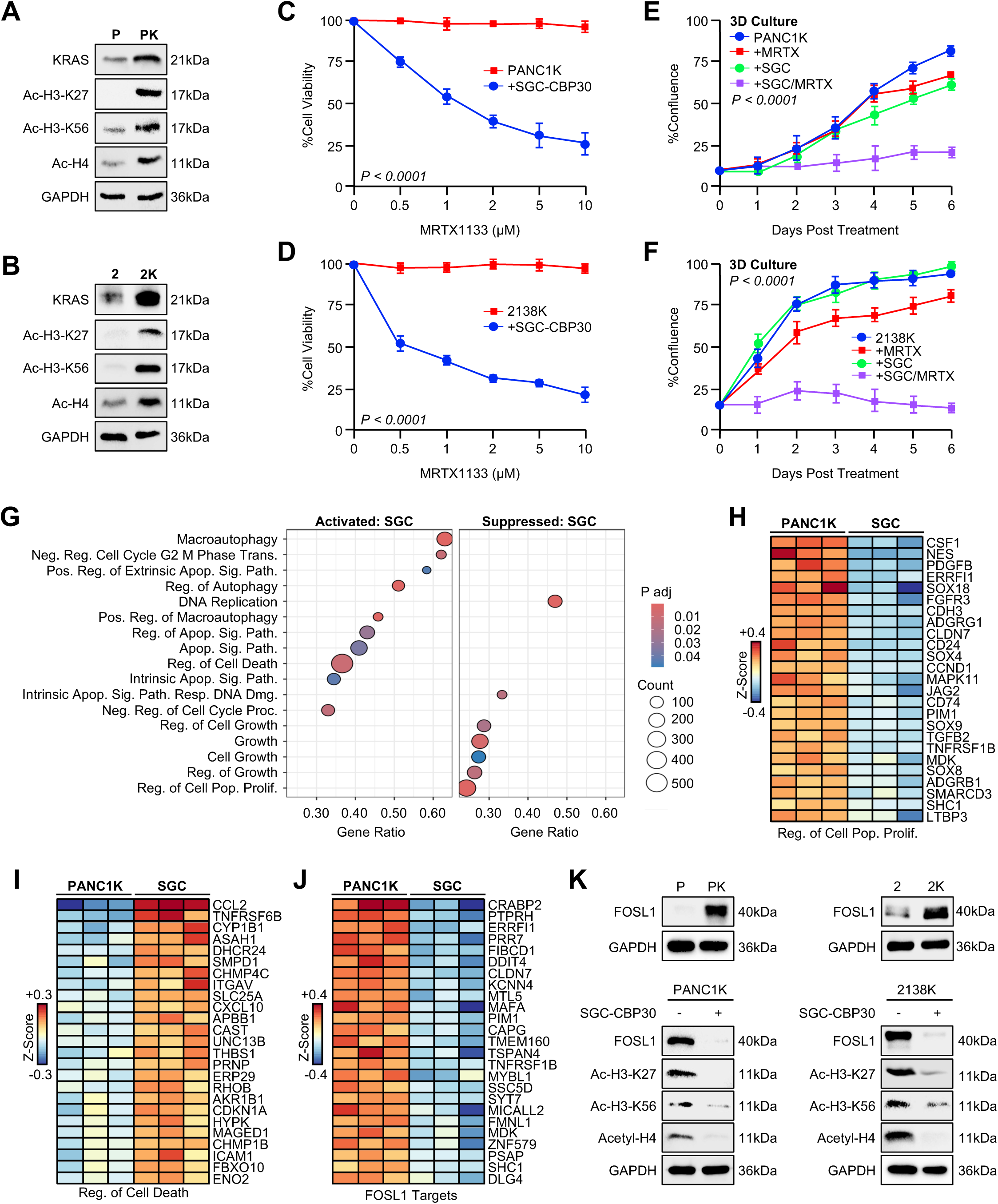
Inhibition of EP300-mediated histone acetylation reverses MRTX1133 resistance and reduces FOSL1 expression **(A,B)** PANC1/PANC1K or 2138/2138K cells were lysed and subjected to western blot for acetylated histones. **(C,D)** PANC1K or 2138K cells were pre-treated with either a DMSO vehicle or 5µM of the EP300 inhibitor SGC-CBP30. After 48 hours, cells were collected in serum-free media with either DMSO or SGC-CBP30 and seeded into 96-well plates (4,000/cells per well). After 24 hours, cells were changed to full-serum media also with either DMSO or SGC-CBP30 and increasing concentrations of MRTX1133. After another 72 hours, cell viability was evaluated by 3-(4,5-Dimethylthiazol-2-yl)-2,5-diphenyltetrazolium bromide (MTT) assay. **(E,F)** PANC1K or 2138K cells were treated with either a DMSO vehicle or 5µM SGC-CBP30 for 72 hours. Cells were then collected and seeded in 3D floating collagen cultures in full serum media with the same treatments, each with and without 0.5µM MRTX1133. Cell confluence was quantified over 6 days. **(G)** PANC1K cells were treated with either a DMSO vehicle or 5µM SGC-CBP30 for 72 hours. Cells were seeded in 3D floating collagen cultures (10,000 cells/well) in full serum media supplemented with 0.5µM MRTX1133 and either a DMSO vehicle or 5µM SGC-CBP30. After 48 hours, cells were subjected to RNA sequencing with gene set enrichment analysis (GSEA). **(H-J)** Focused heatmap is shown for select, significantly altered genes in the regulation of cell population proliferation, growth, and FOSL1 targets gene sets. **(K)** PANC1/PANC1K or 2138/2138K cells were evaluated by western blot for FOSL1 expression. PANC1K or 2138K cells were then treated with 5µM SGC-CBP30 for 72 hours and subjected to western blot for FOSL1 as well as acetylated histones.

Consistent with re-sensitization, in PANC1K or 2138K cells treated with SGC-CBP30, MRTX1133 reduced cell viability in a highly dose-dependent manner (**Figure 2C,D**). MRTX1133-resistant cells were next incubated with either a DMSO vehicle or SGC-CBP30 for 72 hours, after which they were seeded in 3D floating collagen cultures at low cell density and treated with a fixed 0.5 µM dose of MRTX1133 and cell confluence was evaluated over the next six days. Though treatment with SGC-CBP30 alone did not affect the growth of these cells, the addition of SGC-CBP30 restored MRTX1133-induced growth suppression in both PANC1K and 2138K cells (**Figure 2E,F**).

To identify the potential resistance mechanisms being regulated by EP300, we reestablished 3D floating collagen cultures using PANC1K cells treated with either a DMSO vehicle or SGC-CBP30. After 48 hours, cells were subjected to RNA sequencing and GSEA. PANC1K cells treated with SGC-CBP30 had upregulation of several gene sets involved the apoptotic signaling pathways and downregulation of several gene sets involved in cell proliferation (**Figure 2G-I and S1C,D).** SGC-CBP30 treatment also led to the suppression of several known targets of the transcription factor FOSL1 (**Figure 2J**), a central regulator of cell survival that has previously been shown to have a role in resistance to MEK inhibition (37). We next repeated these RNA sequencing experiments using 2138K cells. As previously, treatment with SGC-CBP30 enhanced the expression of gene sets involved in cell death and downregulated several gene sets involved in proliferation and several transcriptional targets of FOSL1 (**Figure S1E-H**).

To determine whether FOSL1 may have a role in the MRTX1133-resistant phenotype, we next compared the relative expression of FOSL1 in both parental and MRTX1133-resistant cell lines by Western blot. Both PANC1K and 2138K cells had increased FOSL1 expression (**Figure 2K**), as well as enrichment of several transcriptional targets of FOSL1 and the related AP-1 complex by RNA sequencing (**Figure S2A-D**). Moreover, when PANC1K or 2138K cells were incubated with SGC-CBP30, we observed reduced FOSL1 expression by Western blot as well as an expected decrease in histone acetylation (**Figure 2K**).

### FOSL1-mediated survival signaling is required for MRTX1133 resistance

To determine whether the observed changes in FOSL1 expression have a role in mediating MRTX1133 resistance, we next generated several additional MRTX1133-resistant PDAC cell lines (**Figure S3**) and incubated each with either a control siRNA (siControl) or siFOSL1. Twenty-four hours after transfection, cells were incubated with increasing concentrations of MRTX1133, and cell viability was evaluated by MTT assay after 72 hours. In each case, the addition of siFOSL1 restored MRTX1133 sensitivity, leading to reduced cell viability in a dose-dependent manner (**Figure 3A,B**). Results were then verified using a second siRNA against FOSL1, showing an identical phenotype (**Figure S4A,B**). Similarly, incubation with siFOSL1 also restored the growth-suppressive effects of MRTX1133 in 3D floating collagen cultures (**Figure S4C,D**). Interestingly, incubating parental cell lines with siFOSL1 did not meaningfully enhance the effects of MRTX1133, suggesting that FOSL1 is largely dispensable for responses to MRTX1133 in naïve cells, but is central to the MRTX1133 resistance (**Figure S4E,F**).

**Figure 3.**
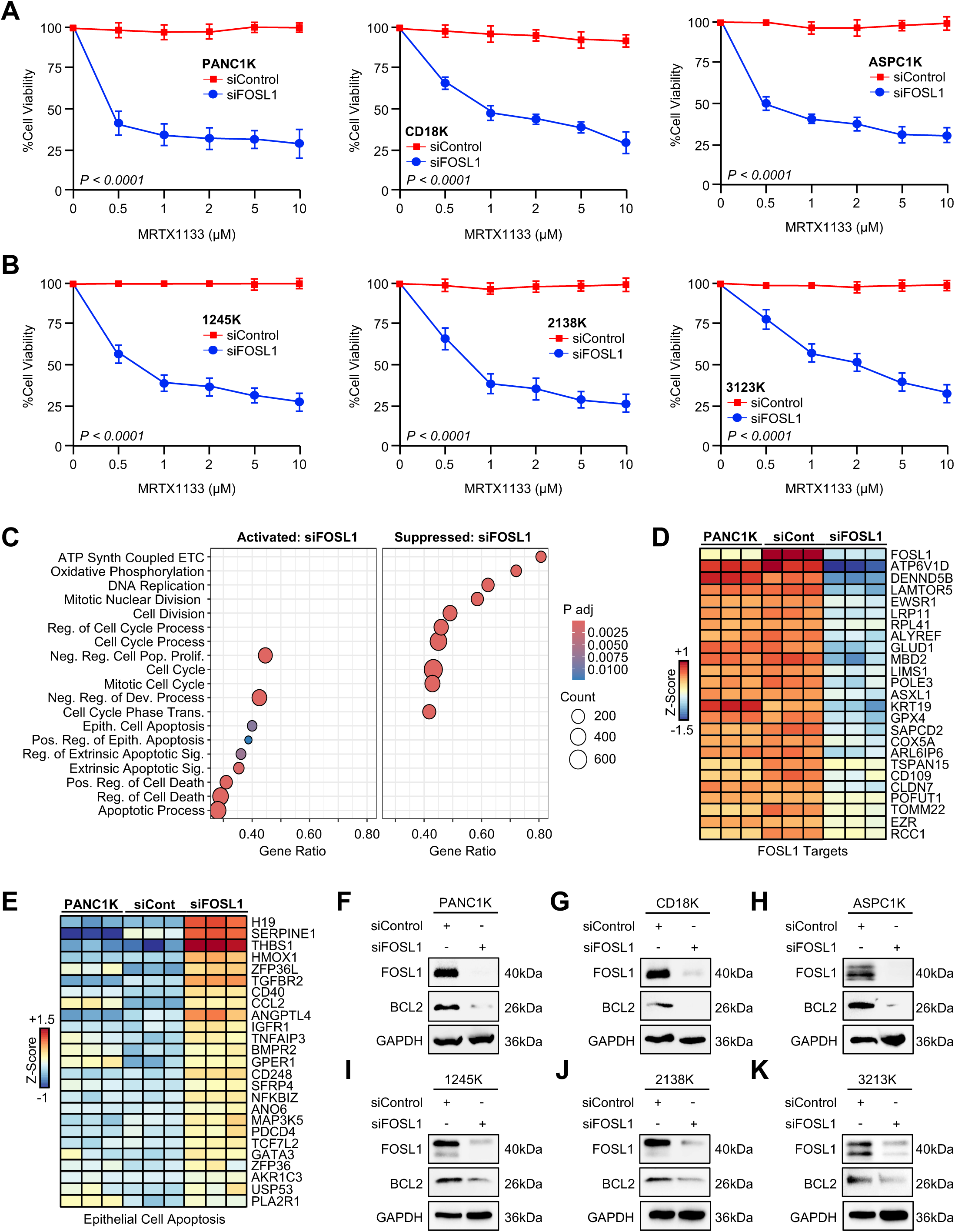
FOSL1-mediated survival signaling is required for MRTX1133 resistance **(A,B)** MRTX1133-resistant cell lines PANC1K, CD18K, ASPC1K, 1245K, 2138K, and 3213K were incubated with either siControl or siFOSL1. After 24 hours, cells were collected, seeded into 96-well plates (4000 cells/well), and incubated with fresh siRNA and increasing concentrations of MRTX1133. Cell viability evaluated by MTT assay after 72 hours. **(C)** PANC1K cells were incubated with either siControl or siFOSL1. After 24 hours, cells were seeded in 3D floating collagen cultures (10,000 cells/well) in full serum media supplemented with fresh siRNA and 0.5µM MRTX1133. After 48 hours, cells were subjected to RNA sequencing with gene set enrichment analysis (GSEA). (**D,E)** Focused heatmap is shown for select, significantly altered genes in the FOSL1 targets and epithelial cell apoptosis gene sets. **(F-K)** MRTX1133-resistant cell lines were serum starved overnight and incubated with either siControl or siFOSL1 in full serum media. After 48 hours, cells were evaluated by western blot for FOSL1 and BCL2 expression.

Given the wide range of FOSL1 targets that could be contributing to the resistant phenotype, we next reestablished 3D floating collagen cultures using PANC1K cells, which had been incubated with either siControl or siFOSL1. After 48 hours, cells were subjected to RNA sequencing and GSEA. Similar to results using SGC-CBP30, PANC1K cells incubated with siFOSL1 demonstrated suppression of several gene sets involved with cell cycle progression and cell proliferation, activation of several gene sets related to cell death (**Figure 3C,D**) and downregulation of known FOSL1 transcriptional targets (**Figure 3E**). Given the observed effects on cell survival signaling, our previous observations linking the cell survival protein BCL2 to MRTX1133 resistance (20), and that BCL2 is a known transcriptional target of FOSL1 (38), drug-resistant cell lines were again incubated with either siControl or siFOSL1 and BCL2 expression evaluated by Western blot. Expectedly, siFOSL1 treatment reduced BCL2 expression in all six MRTX1133-resistant cell lines (**Figure 3F-K**).

### BET inhibitors reverse MRTX1133 resistance and inhibit FOSL1 expression

Given the lack of clinically useful EP300 or FOSL1 inhibitors, we next explored whether BET inhibitors that target acetylation scanning proteins, e.g., BRD4, would similarly reverse the MRTX1133-resistant phenotype, mainly as FOSL1 is a known target of BET inhibitors (39). We, therefore, incubated all six pairs of parental and MRTX1133-resistant PDAC cell lines with either a DMSO vehicle or the BET inhibitors JQ1 or OTX-015, added increasing concentrations of MRTX1133, and evaluated cell viability by MTT assay after 72 hours. Though BET inhibitors did not enhance the responses to MRTX1133 in the parental cell lines (**Figures 4A and S5A**), the addition of BET inhibitors was highly effective in all drug-resistant cell lines, in each case leading to a dose-dependent decrease in cell viability (**Figures 4B and S5B**). We observed similar effects using the additional BET inhibitors ABBV-744 and Trotabresib (**Figure S5C**). Consistent with these observations, the addition of JQ1 or OTX-015 restored MRTX1133-induced growth suppression in both PANC1K and 2138K cells grown in 3D floating collagen cultures (**Figure S5D**).

**Figure 4.**
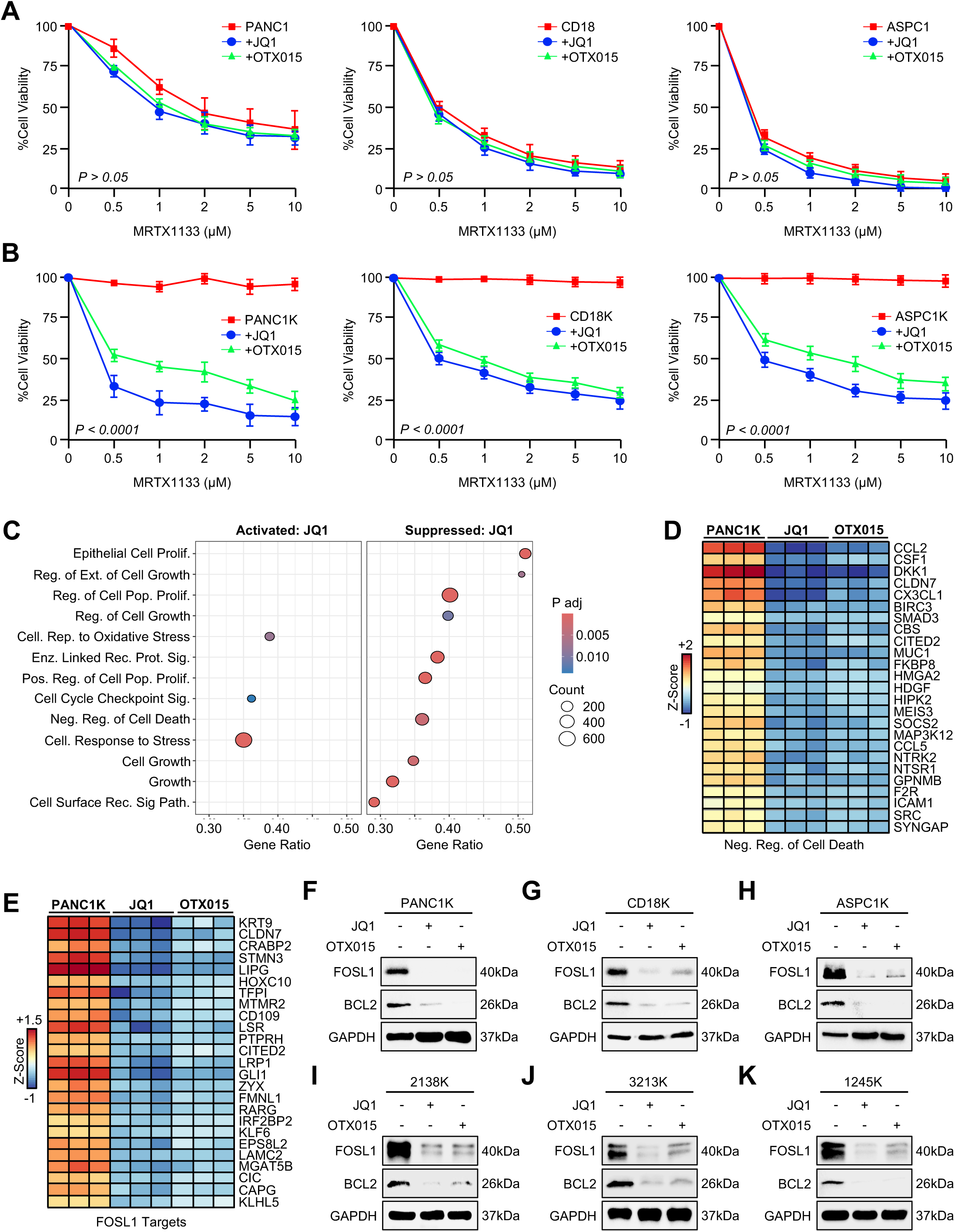
BET inhibitors reverse MRTX1133 resistance and inhibit FOSL1 expression (A,B) Parental PANC1, CD18, or ASPC1 cells were collected in serum-free media and seeded into 96-well plates (4,000/cells per well). Cells were changed to full-serum media containing DMSO vehicle, 1µM JQ1, or 1µM OTX-015 and increasing concentrations of MRTX1133. After another 72 hours, cell viability was evaluated by 3-(4,5-Dimethylthiazol-2-yl)-2,5-diphenyltetrazolium bromide (MTT) assay. This experiment was repeated using the MRTX1133-resistant cell lines PANC1K, CD18K, and ASPC1K. **(C)** PANC1K cells were incubated with either a DMSO vehicle or 1µM JQ1. After 24 hours, cells were seeded in 3D floating collagen cultures in full serum media supplemented with 0.5µM MRTX1133 and either DMSO vehicle or JQ1. After 48 hours, cells were subjected to RNA sequencing with gene set enrichment analysis (GSEA). **(D,E)** Focused heatmap is shown for select, significantly altered genes in the negative regulation of cell death and FOSL1 targets gene sets. **(F-K)** MRTX1133-resistant cell lines were serum starved overnight and incubated with 1µM MRTX1133 and either a DMSO vehicle, 1µM JQ1, or 1µM OTX-015 in full serum media. After 24 hours, cells were evaluated by western blot for FOSL1 and BCL2 expression.

To determine potential mechanisms regulated by BET inhibitors, we again established 3D floating collagen cultures using PANC1K cells, which were then incubated with DMSO vehicle, JQ1, or OTX-015. After 48 hours, cells were subjected to RNA sequencing and GSEA. Similar to incubation with either siFOSL1 or SGC-CBP30, PANC1K cells treated with BET inhibitors had downregulation of several gene sets involved in cell proliferation and cell survival/anti-apoptotic signaling, as well as the upregulation of gene sets involved in cell stress and cell cycle checkpoint signals (**Figure 4C,D and S6A-C**).

BET inhibitors led to the downregulation of FOSL1 and several known transcriptional targets of FOSL1, most notably BCL2 (**Figure 4E and S6D**). Additionally, incubation with either JQ1 or OTX-015 led to the downregulation of FOSL1 itself, as well as its established downstream target BCL2 in all six MRTX1133-resistant PDAC cell lines (**Figure 4F-K**). Consistent with these observations, incubation with the apoptosis inhibitor Q-VD-OPh reversed the tumoricidal effects of both JQ1 and OTX-015 (**Figure S6E**).

### The combination of MRTX1133 and JQ1 improves survival in the orthotopic 1245K model of MRTX1133-resistant PDAC

After demonstrating that MRTX1133-resistant tumor cells maintain their hyperacetylated phenotype in vivo (**Figure S7A,B**), we next sought to evaluate the therapeutic potential of BET inhibition in MRTX1133-resistant PDAC. We first used the 1245K model, which is derived from the well-established FC1245 model developed by the Tuveson group, and utilizes a cell line derived from the transgenic KPC mouse in the C57BL/6 genetic background (40). Per the original reference, when orthotopically injected into the pancreas of a background-matched mouse, this provides a consistent, well-accepted, and physiologically relevant model of PDAC (41). To establish MRTX1133-resistant PDAC tumors, we implanted 10,000 1245K cells into the pancreas of background matched C57BL/6 mice and allowed them to develop ∼250-300 mm^3^ tumors. At this time, mice were randomized into one of four treatment groups at a 50:50 male to female ratio. Mice were treated with daily intraperitoneal injections of DMSO (vehicle control), 30 mg/kg MRTX1133, 50 mg/kg JQ1, or a combination of MRTX1133 and JQ1 (N = 8 mice/group). Mice were euthanized either upon ulceration of the skin or when showing clear signs of health decline, e.g., weight loss, ascites, or lethargy (**Figure 5A,B**).

**Figure 5.**
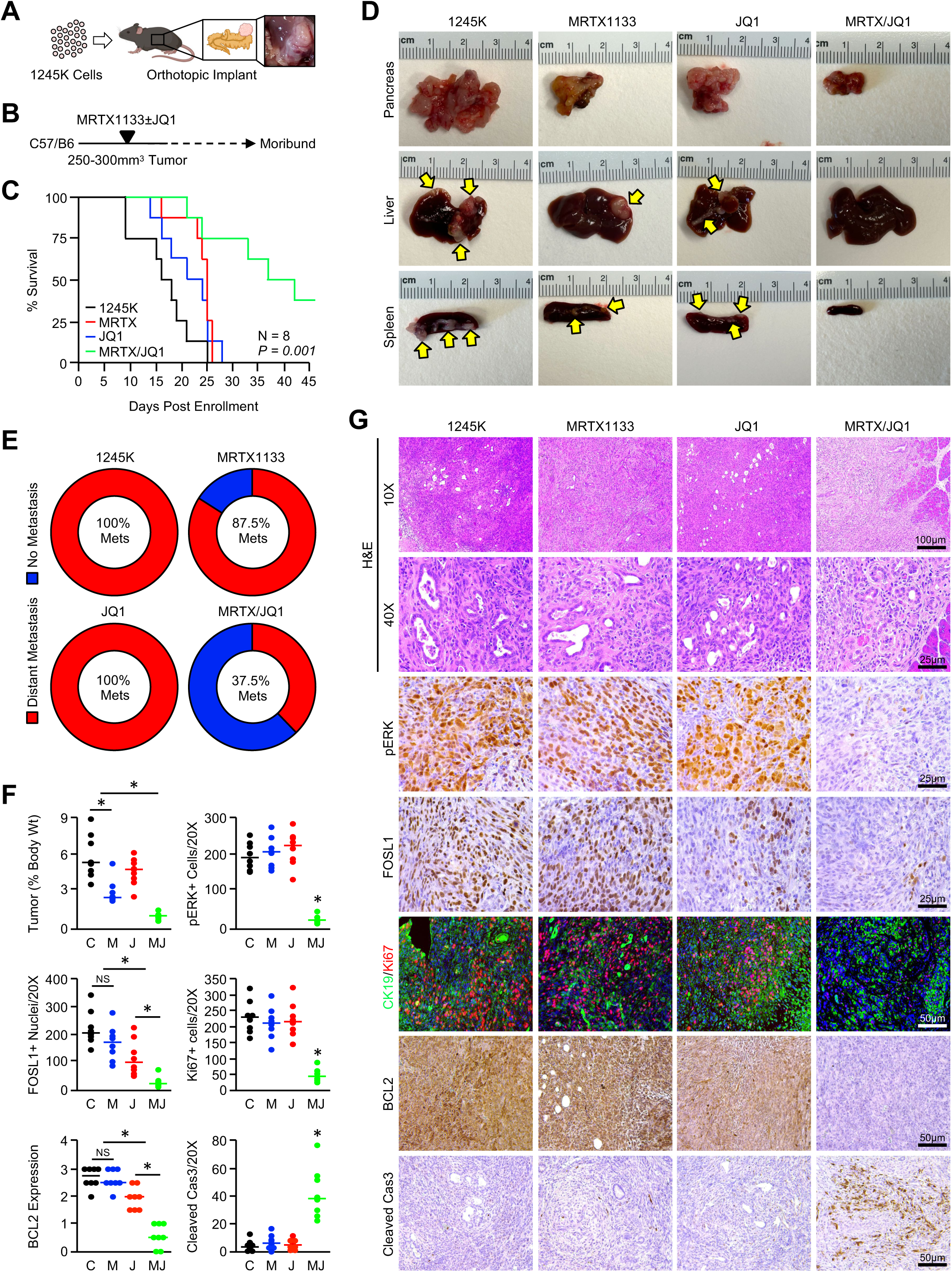
The combination of MRTX1133 and JQ1 improves survival in the orthotopic 1245K model of MRTX1133-resistant PDAC **(A-C)** 1245K cells were collected, resuspended in a 1:1 mixture of Matrigel and full serum media, and 10,000 cells were injected into the pancreas of background matched C57BL/6 mice. Mice were allowed to develop a ∼250-300 mm^3^ tumor, at which point they were enrolled into one of four treatment groups. Mice were treated with daily injections of DMSO (vehicle control), 30 mg/kg MRTX1133, 50 mg/kg JQ1, or a combination of MRTX1133 and JQ1 (N = 8 mice/group). Mice were euthanized either upon ulceration of the skin or when showing clear signs of health decline, e.g., weight loss, ascites, or lethargy. Overall survival is shown via the Kaplan-Meier method. The combination group was significantly different from all other groups, with the P value shown for the combination vs control groups. **(D)** At the study endpoint, gross changes in the pancreas, liver, and spleen were evaluated and representative images shown for each group. **(E)** The percent of mice in each group showing metastatic lesions to the liver or spleen. **(F,G)** Tissues were stained with H&E or via immunohistochemistry for pERK, FOSL1, CK19/Ki67, BCL2, or Cleaved Caspase 3, quantified as described, and results displayed as individual value plots. (C: DMSO Control, M: MRTX1133, J: JQ1, MJ: MRTX1133/JQ1, *p < 0.05).

Tumor-bearing mice treated with MRTX1133 alone had a modest survival advantage compared to controls, as did single-agent treatment with JQ1. However, the combination of MRTX1133 and JQ1 had a significant survival advantage compared with all other groups, with 3/8 mice (37.5%) meeting the survival endpoint compared to 0% in all other groups (**Figure 5C**). All treatments were well tolerated, with no mice showing overt signs of drug-toxicity. On necropsy, 8/8 mice in the control group had extensive nodular masses distributed through a grossly enlarged, firm pancreas (**Figure 5D**). Additionally, 8/8 control mice developed hepatic and/or splenic metastases (**Figure 5D,E**). We observed similar results in the single agent arms, with 8/8 MRTX1133 and 8/8 JQ1-treated mice developing a large, firm, nodular pancreas. However, mice in the MRTX1133 group had a modest reduction in the weight of the pancreas when normalized to body weight (**Figure 5F**). Additionally, 7/8 MRTX1133 and 8/8 JQ1-treated mice had hepatic and/or splenic metastases by the study endpoint (**Figure 5D,E**). In the combination arm, though 5/8 mice had large, frank tumors throughout the pancreas, tumors were generally smaller than those in other groups, and there was a significant reduction in the weight of the pancreas, particularly when normalized to body weight (**Figure 5F**).

On histologic evaluation, mice in the combination arm had some retention of normal gland architecture compared to other groups. Consistent with the drug-resistant phenotype, mice treated with MRTX1133 had no significant reduction in ERK activation, however, mice in the combination group had a significant reduction in staining for activated ERK (**Figure 5F,G**). Mice in the JQ1 group had modestly reduced staining for FOSL1, though mice in the MRTX1133/JQ1 group had more significant suppression of FOSL1 staining. Consistent with the above observations, only mice in the combination group had a decrease in staining for proliferation surrogate Ki67 (**Figure 5F,G**). Additionally, mice in the MRTX1133/JQ1 group had a marked reduction in BCL2 staining with a corresponding increase in apoptosis via cleaved caspase 3 staining (**Figure 5F,G**). We subsequently evaluated tissues from metastatic lesions in the liver, which showed similar changes in ERK activation, FOSL1 expression, proliferation, BCL2 expression, and apoptosis (**Figure S7C,D**).

### The combination of MRTX1133 and JQ1 improves survival in the extremely aggressive 2138K model of MRTX1133-resistant PDAC

To explore the therapeutic efficacy of MRTX1133 and BET inhibition in a more aggressive model system, in parallel studies, we also established tumors using the 2138K syngeneic model, also derived from the transgenic KPC mouse in the C57BL/6 genetic background (20). We implanted 7,500 2138K cells into the pancreas of background-matched C57BL/6 mice and allowed them to develop large ∼800-1,000 mm^3^ tumors. As previously, mice were randomized into one of four treatment groups at a 50:50 male to female ratio. Mice were treated with daily intraperitoneal injections of DMSO (vehicle control), 30 mg/kg MRTX1133, 50 mg/kg JQ1, or a combination of MRTX1133 and JQ1 (N = 10 mice/group). Mice were again euthanized either upon ulceration of the skin or when showing clear signs of health decline, e.g., weight loss, ascites, or lethargy (**Figure 6A,B**).

**Figure 6.**
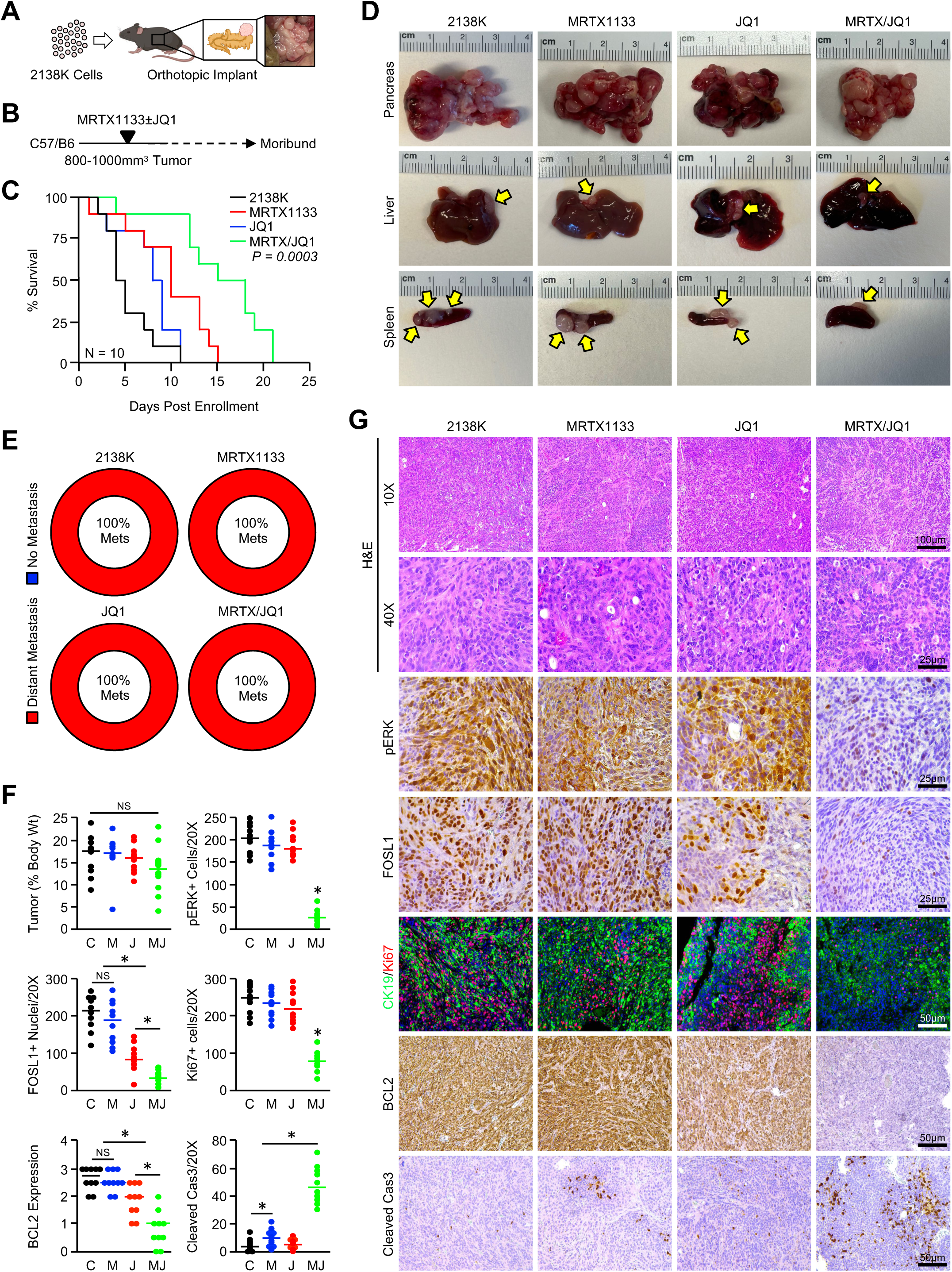
The combination of MRTX1133 and JQ1 improves survival in the extremely aggressive 2138K model of MRTX1133-resistant PDAC **(A-C)** 2138K cells were collected, resuspended in a 1:1 mixture of Matrigel and full serum media, and 7,500 cells were injected into the pancreas of background matched C57BL/6 mice. Mice were allowed to develop a ∼800-1,000 mm^3^ tumor, at which point they were enrolled into one of four treatment groups. Mice were treated with daily injections of DMSO (vehicle control), 30 mg/kg MRTX1133, 50 mg/kg JQ1, or a combination of MRTX1133 and JQ1 (N = 10 mice/group). Mice were euthanized either upon ulceration of the skin or when showing clear signs of health decline, e.g., weight loss, ascites, or lethargy. Overall survival is shown via the Kaplan-Meier method. The combination group was significantly different from all other groups, with the P value shown for the combination vs control groups. **(D)** At the study endpoint, gross changes in the pancreas, liver, and spleen were evaluated and representative images shown for each group. **(E)** The percent of mice in each group showing metastatic lesions to the liver or spleen. **(F,G)** Tissues were stained with H&E or via immunohistochemistry for pERK, FOSL1, CK19/Ki67, BCL2, or Cleaved Caspase 3, quantified as described, and results displayed as individual value plots. (C: DMSO Control, M: MRTX1133, J: JQ1, MJ: MRTX1133/JQ1, *p < 0.05).

Contrasting mice bearing 1245K-derived tumors which had a median survival of 15 days post enrollment (29 days post implantation), mice with 2138K-derived tumors had a median survival of only 4 days post enrollment or two weeks post implantation (**Figure 6C**). Once again, all treatments were well tolerated, with no mice showing overt signs of drug-toxicity. Though treatment with single agent MRTX1133 or JQ1 led to a very modest survival advantage, the combination of MRTX1133 and JQ1 significantly extended overall survival (**Figure 6C**). As previously, all mice in the control or single-agent groups had an enlarged, nodular pancreas. Despite the prolonged survival, at the study endpoint, mice in the combination arm had a similar appearing pancreas to those from all other groups (**Figure 6D**). Consistent with the more aggressive phenotype of the 2138K model, all mice developed splenic or hepatic metastases independent of treatment group (**Figure 6D,E)**. Hence, at the study endpoint, mice in the combination arm had an overall similar tumor burden as those in other groups, only with increased latency most consistent with delayed progression or transient disease stabilization (**Figure 6D-F**).

Consistent with these observations, at their respective endpoints, mice in each group had a similar frequency of tumor nests with no preservation of normal gland architecture. Similar to the 1245K model, Mice treated with JQ1 or MRTX1133/JQ1, mice treated with single agent MRTX1133 again had minimal reduction in ERK activation by immunohistochemistry, though, mice treated with both MRTX1133 and JQ1 had significantly reduced pERK staining (**Figure 6F,G**). Mice in the JQ1 group had some reduced staining in FOSL1, though mice in the MRTX1133/JQ1 group had minimal FOSL1 expression (**Figure 6F,G**). As previously, only mice in the combination group had a decrease in staining for proliferation surrogate Ki67. Though treatment with JQ1 led to a modest decrease in BCL2 staining, mice in the MRTX1133/JQ1 group had a highly significant reduction in BCL2 with was associated with an increase in apoptosis via cleaved caspase 3 staining (**Figure 6F,G**). We similarly evaluated tissues from metastatic lesions in the liver, once again showing similar changes in ERK activation, FOSL1 expression, proliferation, BCL2 expression, and apoptosis (**Figure S8A,B**).

## DISCUSSION

Gain-of-function KRAS mutations are ubiquitous in PDAC tumors, and mutant KRAS has long been proposed as an attractive therapeutic target (4). Recently, KRAS^G12D^ inhibitors have entered the clinical arena, and investigators have been eager to determine their efficacy in PDAC patients (6,7). However, preclinical studies to date suggest that, much like clinical data regarding the use of KRAS^G12C^ inhibitors in PDAC, KRAS^G12D^ inhibitors, including MRTX1133, are likely to provide only transient periods of disease stabilization ultimately followed by progression (9,10). Given the lack of an effective treatment for patients with advanced PDAC, there is an increasing interest in understanding mechanisms of resistance to KRAS^G12D^ inhibitors, as well as identifying effective combination strategies to improve clinical responses or increase latency to progression.

In the present study, we established *in vitro* models of acquired resistance to MRTX1133 and conducted RNA sequencing experiments to evaluate the underlying changes in cell signaling. This approach suggested that the MRTX1133-resistant phenotype is associated a relative increase in histone acetylation, enhancing FOSL1-mediated expression of pro-survival signaling. Accordingly, therapeutic inhibition of the HAT protein EP300, siRNA against FOSL1, or the inhibition of the acetylation scanning BET proteins re-sensitized drug-resistant cell lines to MRTX1133 and impaired the expression of cell survival genes e.g., BCL2. In two MRTX1133-resistant syngeneic tumor models, the combination of MRTX1133 and the BET inhibitor JQ1 markedly enhanced survival, suppressing FOSL1 expression and inducing cell death.

Though the combination of BET and KRAS inhibitors has yet to be explored, there is early preclinical data supporting the combination of BET and MEK inhibitors. For example, MEK inhibitors are often bypassed by rebound upregulation of select receptor tyrosine kinases by *de novo* formation of BRD4-enriched enhancers (43,44). Like our observations, this also appears to be EP300-dependent, as BRD4 and CBP/EP300 inhibition arrested enhancer seeding and compensatory upregulation of receptor tyrosine kinases (43). Accordingly, the combination of JQ1 and the MEK inhibitor Trametinib was highly effective in several PDAC cell lines (45). Similar results have been observed in models of colon cancer, in which JQ1 synergized with Trametinib to induce regression of subcutaneous xenografts (46). This has been observed in other cancer types, as BET inhibition enhanced the effects of Trametinib in xenografted models of glioma (47). Combined BET and MEK inhibition have also been shown to synergize in ovarian tumor models in part by modulating the expression of BCL2 family proteins (48).

Though these and other studies support the combination of RAS pathway and BET inhibition, the mechanisms through which these medications synergize appear to be less understood. FOSL1, also known as FRA1, is emerging as a critical target of KRAS-driven oncogenesis. For instance, a recent study has identified increased FOSL1 expression as a predictor of poor prognosis in KRAS-mutated lung and pancreas cancers (49). Similarly, loss of FOSL1 reduced tumor formation and extended survival in a genetically modified mouse model of advanced lung cancer, and therapeutic inhibition of MEK and the FOSL1-target gene AURKA was highly effective in subcutaneous tumor models (49). A parallel study drew similar conclusions, with FOSL1 deletion inhibiting KRAS-induced lung tumorigenesis, though the authors concluded that these effects were mediated by regulation of the EGF family member Amphiregulin and downstream changes in cell survival proteins (38).

Interestingly, a recent study using genetic-loss-of-function experiments determined that FOSL1 is dispensable for KRAS^G12D^-induced tumorigenesis in murine models of PDAC (37). However, FOSL1 appeared to have a central role in determining responsiveness to MEK inhibition, presumably via adaptive rewiring of oncogenic MEK/ERK signaling. Consistent with these observations, the authors demonstrated that degradation of FOSL1 synergized with MEK inhibition, further supporting FOSL1 signaling as a potential resistance mechanism to inhibition of the MEK/ERK pathway (37). Less is known regarding FOSL1 in the setting of KRAS inhibition, however, a recent CRISPR/Cas9 library screen in the ASPC1 cell line identified FOSL1 as one of several candidate genes that may have a role in therapeutic responses to MRTX1133. The use of siRNA against FOSL1 cooperated with MRTX1133 to suppress the growth of ASPC1 cells *in vitro* (50). However, the combined effect was modest and our observations suggest that FOSL1 is not required for therapeutic responses to MRTX1133 in other sensitive cell lines. As our RNA sequencing data suggested that MRTX1133-resistant tumor cells exhibit marked upregulation of FOSL1 and of its known transcriptional targets, we hypothesized that FOSL1 may be central to the drug-resistant phenotype. Though the effects of FOSL1 on gene expression are highly varied and contextual (51), FOSL1 has been shown to regulate a wide variety of genes with central roles in cell survival. For example, in *KRAS*-mutated lung cancer cells, FOSL1-depletion led to marked downregulation of anti-apoptotic proteins, including BCL2 (38).

Like its upstream regulator FOSL1, BCL2 is emerging as a potential mediator of resistance to KRAS pathway inhibition. For example, the RAS/MEK/ERK pathway has long been known to regulate the expression of BCL2. In PDAC tumor cells, the inhibition of MEK signaling led to the downregulation of BCL2 and other BCL2 family members including BCL-XL and MCL1 (52). Accordingly, the effects of MEK inhibition were potentiated by co-targeting the BCL-2 family proteins in both pancreas and lung cancer cells (53). Similar results have been observed in B-RAF-mutated tumor cell lines, as well as in neuroblastoma cells harboring mutational activation of the RAS/MEK/ERK pathway (54,55). Consistent with these observations, our group has recently demonstrated that therapeutic inhibition of BCL2 with Venetoclax cooperates with MRTX1133 in naïve tumor models, and can re-sensitize drug-resistant tumors to MRTX1133 *in vivo* (20).

Importantly, though we used BCL2 as a surrogate marker given these observations, there are likely several anti-proliferative and anti-apoptotic proteins that are mediating the observed response to BET inhibitors. For example, similar results have been observed regarding the BCL2-family member BCL-XL, as combined BCL-XL and MEK inhibition was highly effective in KRAS-mutated cancer cell lines (56). Similarly, the effects of EGFR inhibitor can also be enhanced by co-targeting BCL2 family members (57). Hence, the observed effects on cell death in the present study are likely not solely limited to BCL2 itself and warrant further exploration in the setting of MRTX1133 resistance.

In summary, we demonstrated that MRTX1133-resistance is associated with aberrant histone acetylation. Inhibition of EP300-mediated acetylation reversed the MRTX1133-resistant phenotype, leading to the suppression of FOSL1-mediated survival signaling, notably downregulation of BCL2. These events were recapitulated using BET inhibitors, which re-sensitized tumor cells to the benefit of MRTX1133 *in vitro* and cooperated with MRTX1133 to extend survival in animal models of MRTX1133-resistant PDAC. As BET inhibitors are under clinical investigation, this may provide for an effective strategy to improve therapeutic responses to MRTX1133 or other selective KRAS inhibitors or re-sensitize tumors that have developed resistance to KRAS inhibition. In addition to the effects on cell survival genes with known roles in mediating resistance to KRAS/MEK inhibitors, e.g., BCL2, this strategy may provide the added benefit of preventing adaptive rewiring of oncogenic MEK/ERK signaling. Hence, the addition of BET inhibitors to selective KRAS inhibitors warrants clinical consideration, particularly once tumors are no longer responsive to KRAS inhibitor monotherapy.

## ACKNOWLEDGEMENTS

This work was supported by NIH F30CA236031 and the Northwestern University Lurie Cancer Center Translational Bridge Award to D.R. Principe, as well as the Robert and Lora Lurie Endowed Professorship, the Harold E. Eisenberg Foundation, the NIH/NCI grant R01CA265997, the Department of Veterans Affairs grant I01BX005595 to H.G. Munshi, and NIH/NCI training grant T32CA268935 to A.M. Marinelarena. Parts of this work were performed at the Robert H. Lurie Comprehensive Cancer Center (RHLCCC) Pathology Core at Northwestern University, which is supported by the NCI CCSG P30 CA060553 awarded to the RHLCCC. The funding agencies had no role in the design of the study; the collection, analysis, or interpretation of the data; the writing of the manuscript; or the decision to submit the manuscript for publication.

## Author contributions

- DRP: Conceptualization, Formal Analysis, Investigation, Visualization, Writing-original draft, Writing-review and editing
- JHB: Investigation
- AEM: Investigation
- AMM: Investigation
- TDP: Investigation
- AFA: Investigation, Formal Analysis
- HGM: Conceptualization, Formal Analysis, Writing-review and editing

