## Supplemental Materials for "Resistance to the KRAS^G12D^ Inhibitor MRTX1133 is Associated with Increased Sensitivity to BET Inhibition"

[1]Department of Medicine, Physician Scientist Training Program, University of Wisconsin, Madison, WI; [2]Department of Medicine, Feinberg School of Medicine, Northwestern University, Chicago, IL; [3]Jesse Brown VA Medical Center, Chicago, IL; [4]The Robert H. Lurie Comprehensive Cancer Center, Chicago, IL; [5]Division of Genetics, Department of Morphology and Genetics, Federal University of São Paulo, São Paulo, Brazil.

Short Title: MRTX1133-Resistant PDAC Tumors are Sensitive to BET Inhibitors

| | |
|---|---|
| Pages | 7 |
| Tables | 1 |
| Figures | 8 |
| Words | 1,479 |

*Correspondence to:

Daniel R. Principe MD, PhD
Department of Medicine
University of Wisconsin
1685 Highland Avenue
Madison, WI 53705, USA


or

Hidayatullah G. Munshi MD
Department of Medicine
Northwestern University Feinberg School of Medicine
303 E. Superior Ave., Lurie 3-117
Chicago, IL 60611, USA


**Conflict of Interest Disclosure:** The authors have no conflicts to disclose.

#### SUPPLEMENTAL TABLE

| Antibody | Application | Vendor | Product Number | Dilution |
|---|---|---|---|---|
| Ac-H3-K9 | WB | Cell Signaling | 9649S | 1:1000 |
| Ac-H3-K27 | WB | Millipore | 05-1334 | 1:1000 |
| Ac-H3-K56 | WB | Millipore | 04-1135 | 1:1000 |
| Ac-H4 | WB/IHC | Millipore | 07-329 | 1:1000 |
| BCL2 (Human) | WB | Cell Signaling | 3498S | 1:1000 |
| BCL2 (Mouse) | WB | BD Biosciences | 554218 | 1:1000 |
| BCL2 (Mouse) | IHC | abcam | AB182858 | 1:500 |
| CK19 | IF | University of Iowa Hybridoma Bank | TROMA-III | 1:100 |
| Cleaved Cas3 | IHC | Cell Signaling | 9664L | 1:1000 |
| ERK | WB | Cell Signaling | 9102S | 1:1000 |
| FOSL1 (Human) | WB | Cell Signaling | 5281S | 1:1000 |
| FOSL1 (Mouse) | WB | abcam | AB252421 | 1:1000 |
| FOSL1 (Mouse) | IHC | Cell Signaling | 28801S | 1:100 |
| GAPDH | WB | EMD Millipore | MAB374 | 1:2000 |
| Ki67 | IHC/IF | Cell Signaling | 12202S | 1:1000 |
| KRAS | WB | Novus Biologicals | NBP2-45536 | 1:2000 |
| Pan Histone | WB | Millipore | MABE71 | 1:1000 |
| pERK | WB | Cell Signaling | 9101S | 1:1000 |
| pERK | IHC | Cell Signaling | 9101S | 1:500 |
| pMEK | WB | Cell Signaling | 9154S | 1:1000 |
| MEK | WB | Cell Signaling | 9122S | 1:1000 |

**Table S1. Antibodies arranged by vendor**

#### SUPPLEMENTAL FIGURE LEGENDS

##### Figure S1. Inhibition of EP300-mediated histone acetylation enhances apoptotic signaling in MRTX1133-resistant PDAC cells

**(A)** Parental and KRAS-inhibitor resistant cells were serum-starved overnight, pre-treated with 2µM MRTX1133 for 30 minutes, after which they were changed to full serum media containing 2µM MRTX1133. After 24 hours, pMEK was evaluated by western blot. **(B)** PANC1/PANC1K or 2138/2138K cells were lysed and subjected to western blot for total histones. **(C,D)** PANC1K cells were treated with DMSO (vehicle control) or 5µM SGC-CBP30 for 72 hours. Cells were seeded in 3D floating collagen cultures (10,000 cells/well) in full serum media supplemented with 0.5µM MRTX1133 and DMSO or 5µM SGC-CBP30. After 48 hours, cells were subjected to RNA sequencing. Focused heatmap is shown for select, significantly altered genes in the growth and intrinsic apoptotic gene sets. **(E)** 2138K cells were treated with either a DMSO vehicle or 5µM SGC-CBP30 for 72 hours. Cells were seeded in 3D floating collagen cultures (10,000 cells/well) in full serum media supplemented with 0.5µM MRTX1133 and either a DMSO vehicle or 5µM SGC-CBP30. After 48 hours, cells were subjected to RNA sequencing with gene set enrichment analysis (GSEA). **(F-H)** Focused heatmap is shown for select, significantly altered genes in the regulation of cell population proliferation, negative regulation of cell death, and FOSL1 targets gene sets.

##### Figure S2. MRTX1133 resistance is associated increased FOSL1/AP-1 signaling

**(A)** PANC1/PANC1K or **(B)** 2138/2138K tumor cells were seeded in 3D floating collagen cultures for 48 hours and subjected to RNA sequencing and gene set enrichment analysis (GSEA). Focused heatmap is shown for select, significantly altered genes in the FOSL1 targets gene set. **(C,D)** Focused heatmap is shown for select, significantly altered genes in the AP-1 targets gene set.

**Figure S3. Validation of additional *in vitro* models of MRTX1133 resistance**

**(A-D)** Murine 3213 and 1245 or human CD18 and ASPC1 cancer cell lines were incubated with increasing concentrations of MRTX1133 until viable in 2μM. After this point, cells were referred as 3213K, 1245K, CD18K, or ASPC1K for KRAS-inhibitor resistant cells. Cells were collected in serum-free media and seeded into 96-well plates (4,000/cells per well). Cells were changed to full-serum media containing either a DMSO vehicle or increasing concentrations of MRTX1133. After another 72 hours, cell viability was evaluated by 3-(4,5-Dimethylthiazol-2-yl)-2,5-diphenyltetrazolium bromide (MTT) assay.

**Figure S4. Loss of FOSL1 reverses MRTX1133 resistance in 3D floating collagen cultures**

**(A,B)** MRTX1133-resistant cell lines PANC1K and 2138K cells were incubated with either siControl or a second siRNA against FOSL1 (siFOSL1 #2). After 24 hours, cells were collected, seeded into 96-well plates (4000 cells/well), and incubated with fresh siRNA and increasing concentrations of MRTX1133. Cell viability was evaluated by MTT assay after 72 hours. **(C,D)** PANC1K or 2138K cells were incubated with either siControl or siFOSL1. After 24 hours, cells were seeded in 3D floating collagen cultures (10,000 cells/well) in full serum media supplemented with fresh siRNA and 0.5μM MRTX1133. Cell confluence was quantified, and growth shown over a period of 5 days. **(E,F)** MRTX1133-naïve cell lines PANC1 and 2138 cells were incubated with either siControl or one of two siRNAs against FOSL1. After 24 hours, cells were collected, seeded into 96-well plates (4000 cells/well), and incubated with fresh siRNA and increasing concentrations of MRTX1133. Cell viability was evaluated by MTT assay after 72 hours.

**Figure S5. BET inhibitors reverse MRTX1133 resistance in additional PDAC cell lines**

**(A)** Parental 2138, 3213, and 1245 murine PDAC cell lines were collected in serum-free media and seeded into 96-well plates (4,000/cells per well). Cells were changed to full-serum media containing DMSO, 1μM JQ1, or 1μM OTX-015 and increasing concentrations of MRTX1133. After another 72 hours, cell viability was evaluated by 3-(4,5-Dimethylthiazol-2-yl)-2,5-diphenyltetrazolium bromide (MTT) assay. **(B)** This experiment was repeated using the MRTX1133-resistant 2138K, 3213K, and 1245K cell lines. **(C)** PANC1K or 2138K cells were collected in serum-free media seeded into 96-well plates (4,000/cells per well). Cells were changed to full-serum media containing DMSO, 1μM ABBV-744, or 1μM Trotabresib and increasing concentrations of MRTX1133. **(D)** PANC1K or 2138K cells were incubated with either a DMSO vehicle, 1μM JQ1, or 1μM OTX-015. After 24 hours, cells were seeded in 3D floating collagen cultures (10,000 cells/well) and incubated with either a DMSO vehicle, a fixed 0.5 μM dose of MRTX1133, JQ1, OTX-015, JQ1/MRTX1133, or OTX-015/MRTX1133. Cell confluence was quantified, and growth shown over a period of 6 days.

**Figure S6. BET inhibitors reverse MRTX1133 resistance in 3D floating collagen cultures**

**(A)** PANC1K cells were incubated with either a DMSO vehicle or 1μM JQ1. After 24 hours, cells were seeded in 3D floating collagen cultures in full serum media supplemented with 0.5μM MRTX1133 and either DMSO vehicle, JQ1, or 1μM OTX-015. After 48 hours, cells were subjected to RNA sequencing with gene set enrichment analysis (GSEA). Results are shown for DMSO and OTX-015-treated cells. For JQ1 data see Figure 4C. **(B,C)** Focused heatmap is shown for select, significantly altered genes in epithelial cell proliferation and cell cycle checkpoint gene sets. **(D)** PANC1K or 2138K were incubated with either a DMSO vehicle, 1μM JQ1, or 1μM OTX-015. After 48 hours, mRNA expression of FOSL1 and BCL2 were evaluated by qPCR. **(E)** PANC1K or 2138K cells were collected in serum-free media and next seeded into 96-well plates (4,000/cells per well). Cells were changed to full-serum media containing DMSO vehicle, 1μM JQ1, or 1μM OTX-015, Q-VD-Oph (Q-VD), JQ1/Q-VD, or OTX-015/Q-VD, each with increasing concentrations of MRTX1133. After another 72 hours, cell viability was evaluated by 3-(4,5-Dimethylthiazol-2-yl)-2,5-diphenyltetrazolium bromide (MTT) assay.

**Figure S7. The combination of MRTX1133 and JQ1 reduces FOSL1 and BCL2 expression in metastatic lesions in the 1245K tumor model**

**(A,B)** 1245, 1245K, 2138, or 2138K cells were collected, resuspended in a 1:1 mixture of Matrigel and full serum media, and 25,000 cells were injected into the flanks of background matched C57BL/6 mice. Mice were euthanized when tumor volume exceeded 1200-1500 $mm^3$, ulcerated, or when mice showed clear signs of health decline, e.g., weight loss, ascites, or lethargy. At the study endpoint, tissues were collected and stained with H&E or via immunohistochemistry for acetyl-H4, quantified as described, and results displayed as individual value plots. **(C,D)** 1245K cells were collected, resuspended in a 1:1 mixture of Matrigel and full serum media, and 10,000 cells were injected into the pancreas of background matched C57BL/6 mice. Mice were allowed to develop a ~250-300 $mm^3$ tumor, at which point they were enrolled into one of four treatment groups. Mice were treated with daily injections of DMSO (vehicle control), 30 mg/kg MRTX1133, 50 mg/kg JQ1, or a combination of MRTX1133 and JQ1 (N = 8 mice/group). Mice were euthanized either upon ulceration of the skin or when showing clear signs of health decline, e.g., weight loss, ascites, or lethargy. At the study endpoint, metastatic lesions from the liver were sectioned and stained with H&E or via immunohistochemistry for pERK, FOSL1, Ki67, BCL2, or Cleaved Caspase 3, quantified as described, and results displayed as individual value plots. (C: DMSO Control, M: MRTX1133, J: JQ1, MJ: MRTX1133/JQ1, *$p < 0.05$).

**Figure S8. The combination of MRTX1133 and JQ1 reduces FOSL1 and BCL2 expression in metastatic lesions in the 2138K tumor model**

**(A,B)** 2138K cells were collected, resuspended in a 1:1 mixture of matrigel and full serum media, and 7,500 cells injected were into the pancreas of background matched C57BL/6 mice. Mice were allowed to develop a ~800-1,000 $mm^3$ tumor, at which point they were enrolled into one of four treatment groups. Mice were treated with daily injections of DMSO (vehicle control), 30 mg/kg MRTX1133, 50 mg/kg JQ1, or a combination of MRTX1133 and JQ1 (N = 10 mice/group). Mice were euthanized either upon ulceration of the skin or when showing clear signs of health decline, e.g., weight loss, ascites, or lethargy. At the study endpoint, metastatic lesions from the liver were sectioned and stained with H&E or via immunohistochemistry for pERK, FOSL1, Ki67, BCL2, or Cleaved Caspase 3, quantified as described, and results displayed as individual value plots. (C: DMSO Control, M: MRTX1133, J: JQ1, MJ: MRTX1133/JQ1, *$p < 0.05$).

### Figure S1

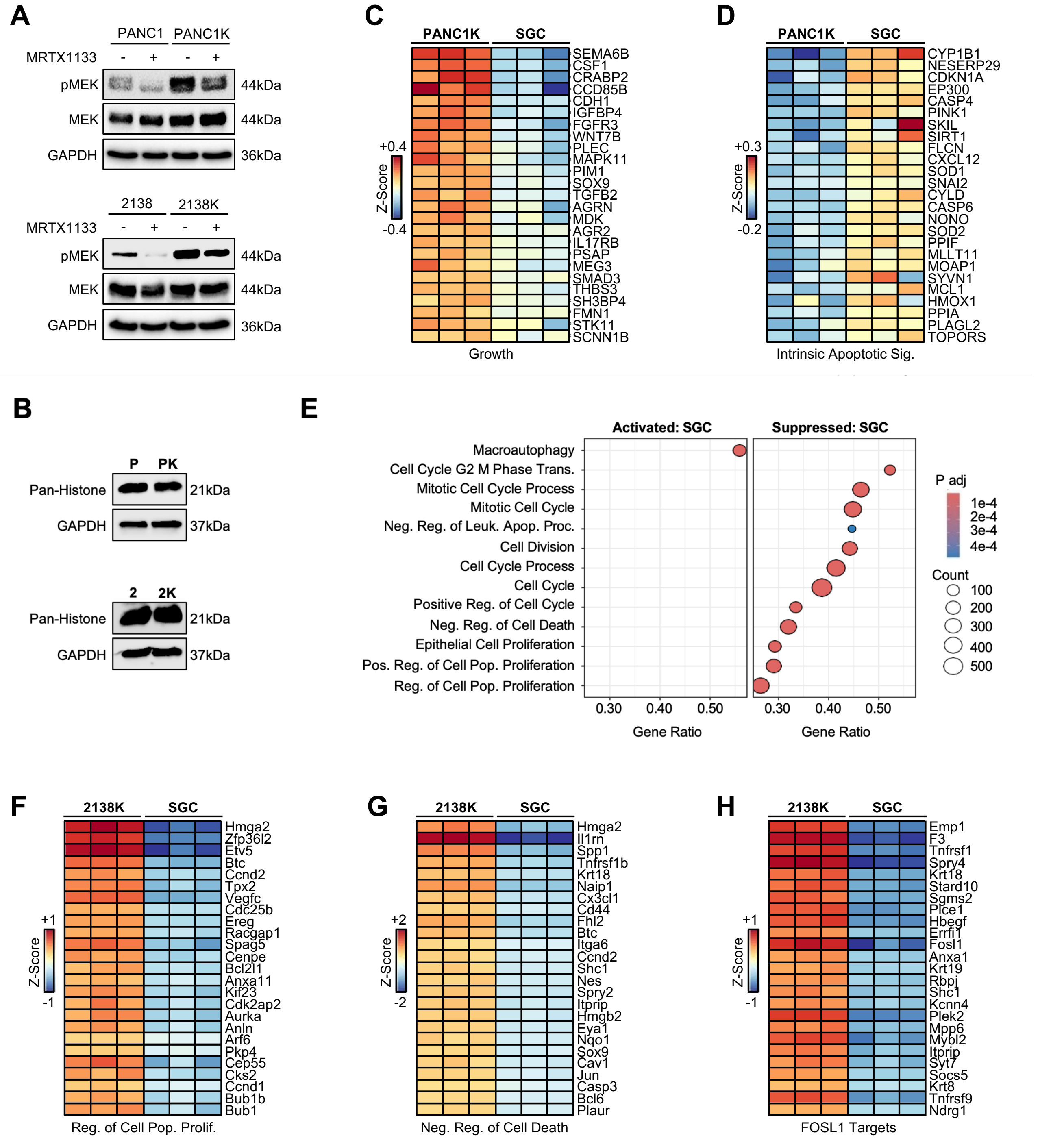


**Figure S1. Inhibition of EP300-mediated histone acetylation enhances apoptotic signaling in MRTX1133-resistant PDAC cells**

**(A)** Parental and KRAS-inhibitor resistant cells were serum-starved overnight, pre-treated with 2μM MRTX1133 for 30 minutes, after which they were changed to full serum media containing 2μM MRTX1133. After 24 hours, pMEK was evaluated by western blot. **(B)** PANC1/PANC1K or 2138/2138K cells were lysed and subjected to western blot for total histones. **(C,D)** PANC1K cells were treated with DMSO (vehicle control) or 5μM SGC-CBP30 for 72 hours. Cells were seeded in 3D floating collagen cultures (10,000 cells/well) in full serum media supplemented with 0.5μM MRTX1133 and DMSO or 5μM SGC-CBP30. After 48 hours, cells were subjected to RNA sequencing. Focused heatmap is shown for select, significantly altered genes in the growth and intrinsic apoptotic gene sets. **(E)** 2138K cells were treated with either a DMSO vehicle or 5μM SGC-CBP30 for 72 hours. Cells were seeded in 3D floating collagen cultures (10,000 cells/well) in full serum media supplemented with 0.5μM MRTX1133 and either a DMSO vehicle or 5μM SGC-CBP30. After 48 hours, cells were subjected to RNA sequencing with gene set enrichment analysis (GSEA). **(F-H)** Focused heatmap is shown for select, significantly altered genes in the regulation of cell population proliferation, negative regulation of cell death, and FOSL1 targets gene sets.

### Figure S2

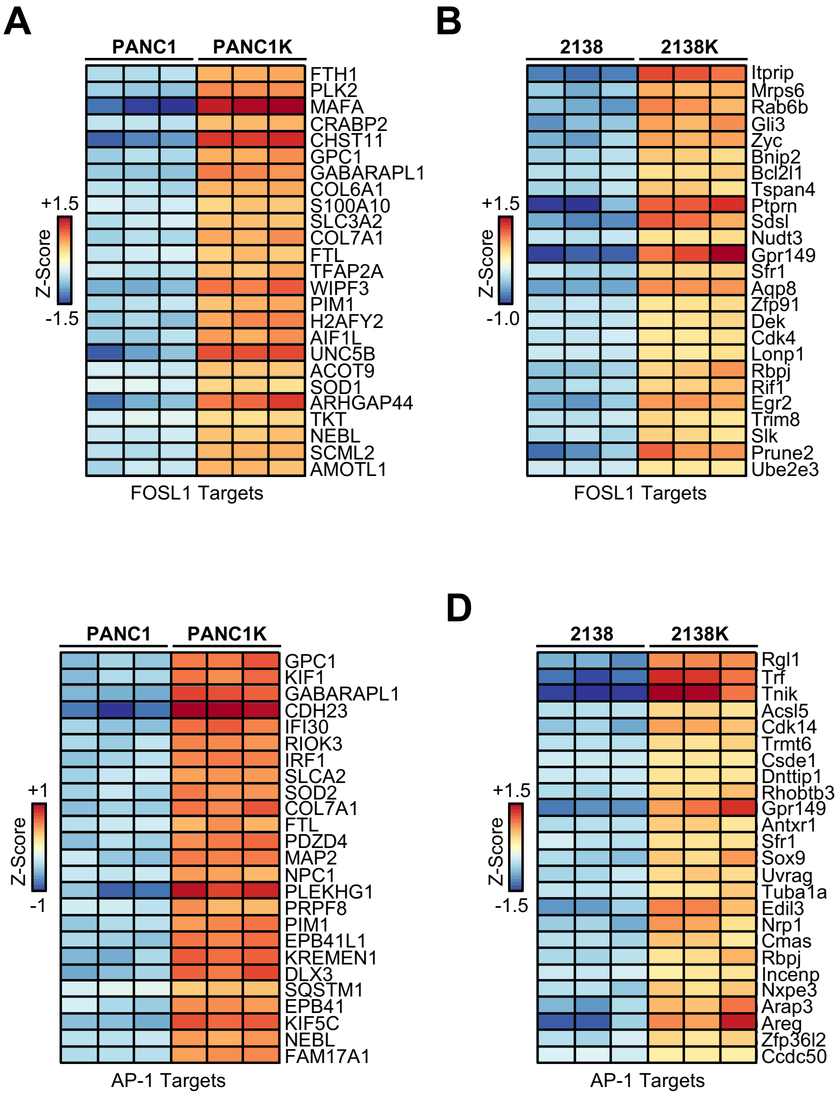


**Figure S2. MRTX1133 resistance is associated increased FOSL1/AP-1 signaling**
**(A)** PANC1/PANC1K or **(B)** 2138/2138K tumor cells were seeded in 3D floating collagen cultures for 48 hours and subjected to RNA sequencing and gene set enrichment analysis (GSEA). Focused heatmap is shown for select, significantly altered genes in the FOSL1 targets gene set. **(C,D)** Focused heatmap is shown for select, significantly altered genes in the AP-1 targets gene set.

### Figure S3

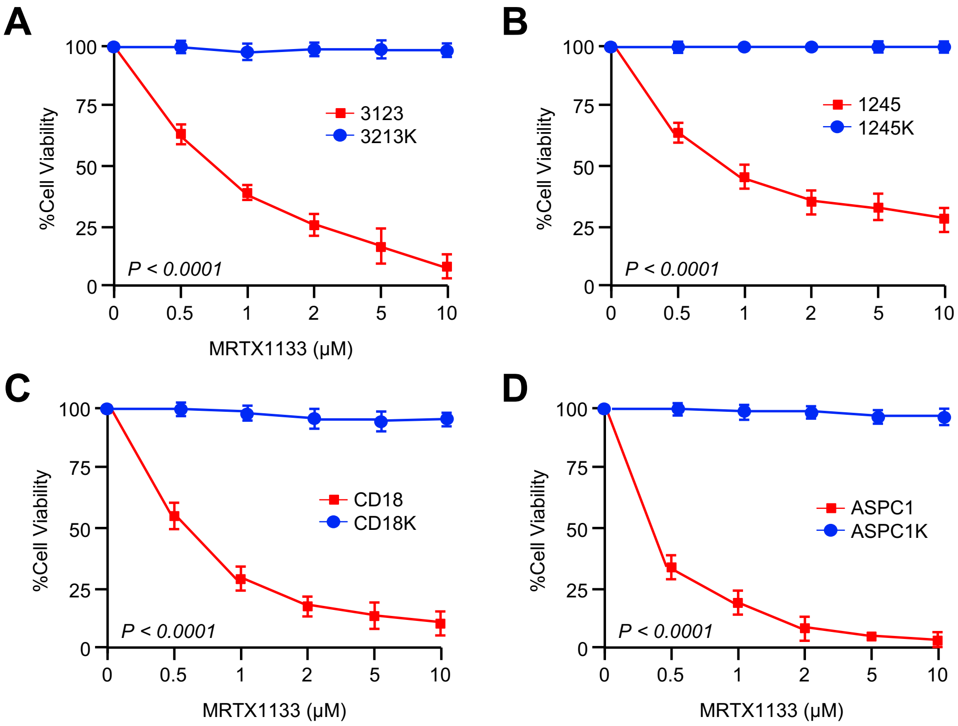


**Figure S3. Validation of additional *in vitro* models of MRTX1133 resistance**

**(A-D)** Murine 3213 and 1245 or human CD18 and ASPC1 cancer cell lines were incubated with increasing concentrations of MRTX1133 until viable in 2μM. After this point, cells were referred as 3213K, 1245K, CD18K, or ASPC1K for KRAS-inhibitor resistant cells. Cells were collected in serum-free media and seeded into 96-well plates (4,000/cells per well). Cells were changed to full-serum media containing either a DMSO vehicle or increasing concentrations of MRTX1133. After another 72 hours, cell viability was evaluated by 3-(4,5-Dimethylthiazol-2-yl)-2,5-diphenyltetrazolium bromide (MTT) assay.

### Figure S4

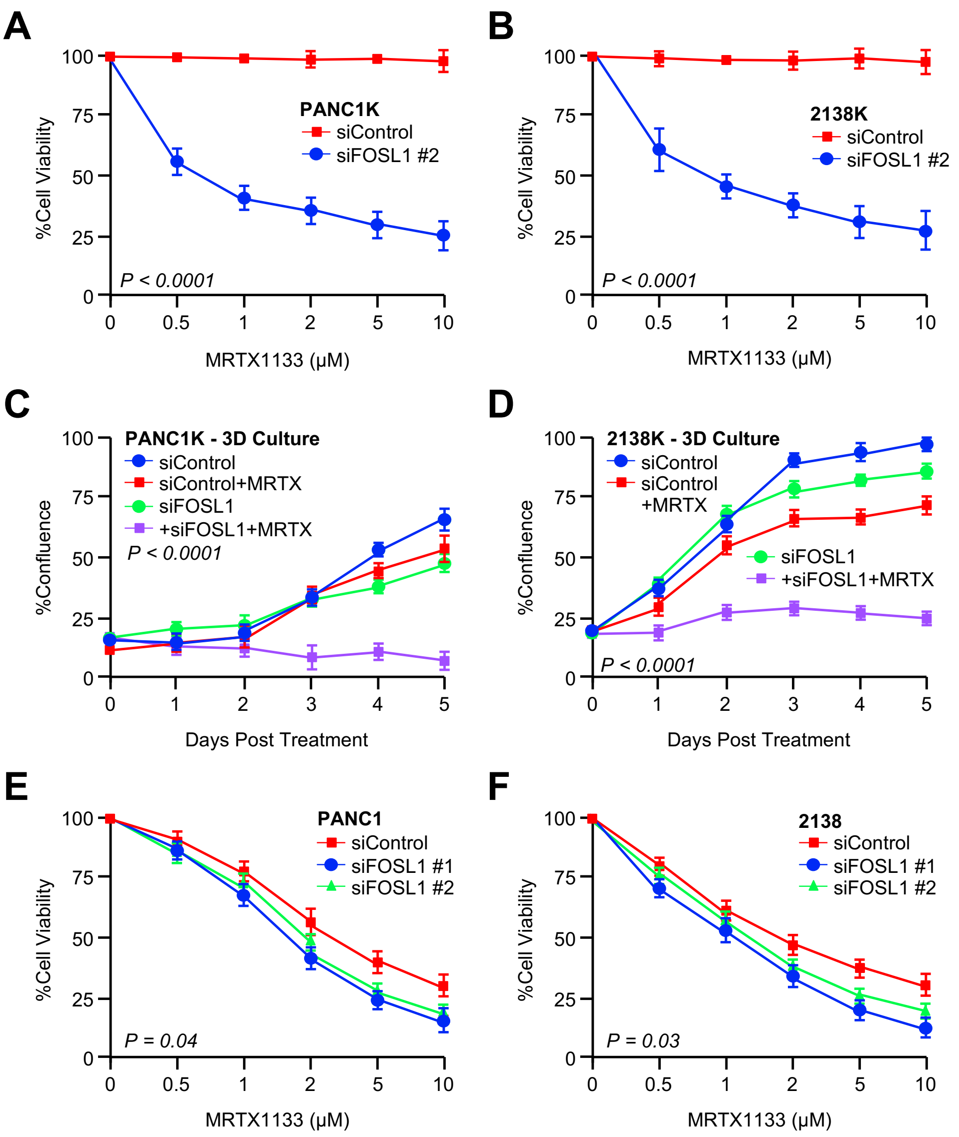


**Figure S4. Loss of FOSL1 reverses MRTX1133 resistance in 3D floating collagen cultures**
**(A,B)** MRTX1133-resistant cell lines PANC1K and 2138K cells were incubated with either siControl or a second siRNA against FOSL1 (siFOSL1 #2). After 24 hours, cells were collected, seeded into 96-well plates (4000 cells/well), and incubated with fresh siRNA and increasing concentrations of MRTX1133. Cell viability was evaluated by MTT assay after 72 hours. **(C,D)** PANC1K or 2138K cells were incubated with either siControl or siFOSL1. After 24 hours, cells were seeded in 3D floating collagen cultures (10,000 cells/well) in full serum media supplemented with fresh siRNA and 0.5µM MRTX1133. Cell confluence was quantified, and growth shown over a period of 5 days. **(E,F)** MRTX1133-naïve cell lines PANC1 and 2138 cells were incubated with either siControl or one of two siRNAs against FOSL1. After 24 hours, cells were collected, seeded into 96-well plates (4000 cells/well), and incubated with fresh siRNA and increasing concentrations of MRTX1133. Cell viability was evaluated by MTT assay after 72 hours.

### Figure S5

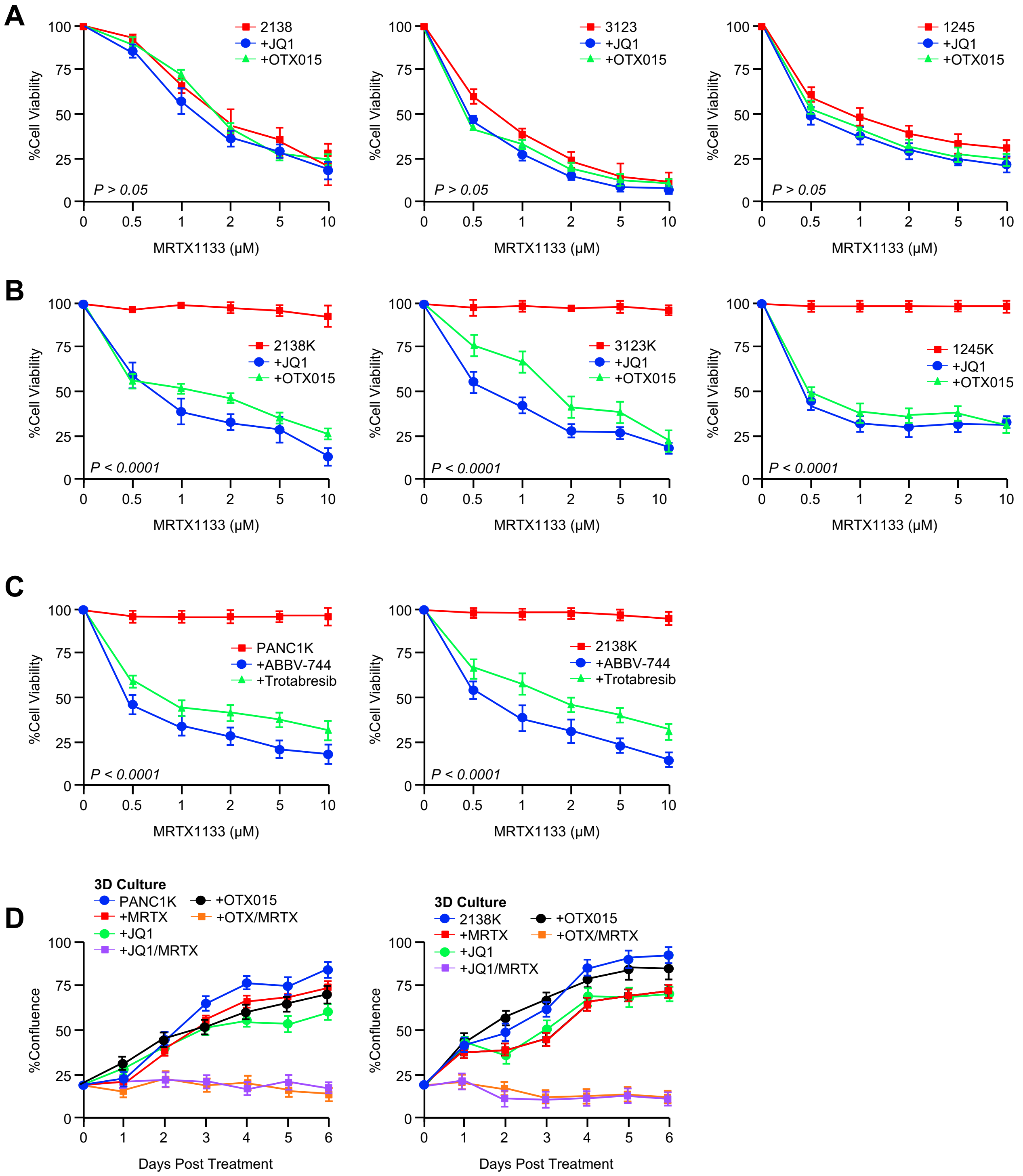


**Figure S5. BET inhibitors reverse MRTX1133 resistance in additional PDAC cell lines**
**(A)** Parental 2138, 3213, and 1245 murine PDAC cell lines were collected in serum-free media and seeded into 96-well plates (4,000/cells per well). Cells were changed to full-serum media containing DMSO, 1μM JQ1, or 1μM OTX-015 and increasing concentrations of MRTX1133. After another 72 hours, cell viability was evaluated by 3-(4,5-Dimethylthiazol-2-yl)-2,5-diphenyltetrazolium bromide (MTT) assay. **(B)** This experiment was repeated using the MRTX1133-resistant 2138K, 3213K, and 1245K cell lines. **(C)** PANC1K or 2138K cells were collected in serum-free media seeded into 96-well plates (4,000/cells per well). Cells were changed to full-serum media containing DMSO, 1μM ABBV-744, or 1μM Trotabresib and increasing concentrations of MRTX1133. **(D)** PANC1K or 2138K cells were incubated with either a DMSO vehicle, 1μM JQ1, or 1μM OTX-015. After 24 hours, cells were seeded in 3D floating collagen cultures (10,000 cells/well) and incubated with either a DMSO vehicle, a fixed 0.5 μM dose of MRTX1133, JQ1, OTX-015, JQ1/MRTX1133, or OTX-015/MRTX1133. Cell confluence was quantified, and growth shown over a period of 6 days.

### Figure S6

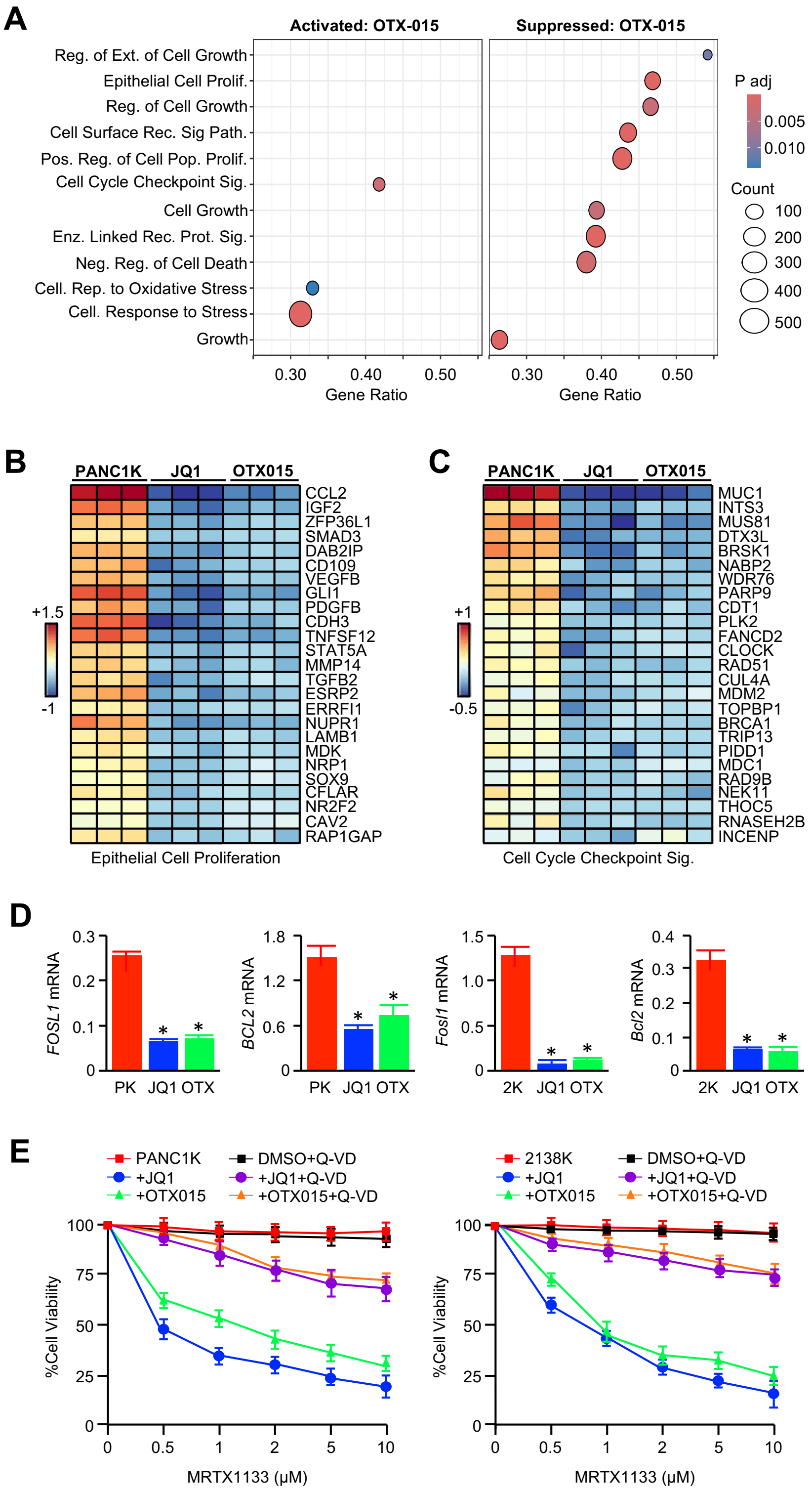


**Figure S6. BET inhibitors reverse MRTX1133 resistance in 3D floating collagen cultures**
**(A)** PANC1K cells were incubated with either a DMSO vehicle or 1μM JQ1. After 24 hours, cells were seeded in 3D floating collagen cultures in full serum media supplemented with 0.5μM MRTX1133 and either DMSO vehicle, JQ1, or 1μM OTX-015. After 48 hours, cells were subjected to RNA sequencing with gene set enrichment analysis (GSEA). Results are shown for DMSO and OTX-015-treated cells. For JQ1 data see Figure 4C. **(B,C)** Focused heatmap is shown for select, significantly altered genes in epithelial cell proliferation and cell cycle checkpoint gene sets. **(D)** PANC1K or 2138K were incubated with either a DMSO vehicle, 1μM JQ1, or 1μM OTX-015. After 48 hours, mRNA expression of FOSL1 and BCL2 were evaluated by qPCR. **(E)** PANC1K or 2138K cells were collected in serum-free media and next seeded into 96-well plates (4,000/cells per well). Cells were changed to full-serum media containing DMSO vehicle, 1μM JQ1, or 1μM OTX-015, Q-VD-Oph (Q-VD), JQ1/Q-VD, or OTX-015/Q-VD, each with increasing concentrations of MRTX1133. After another 72 hours, cell viability was evaluated by 3-(4,5-Dimethylthiazol-2-yl)-2,5-diphenyltetrazolium bromide (MTT) assay.

### Figure S7

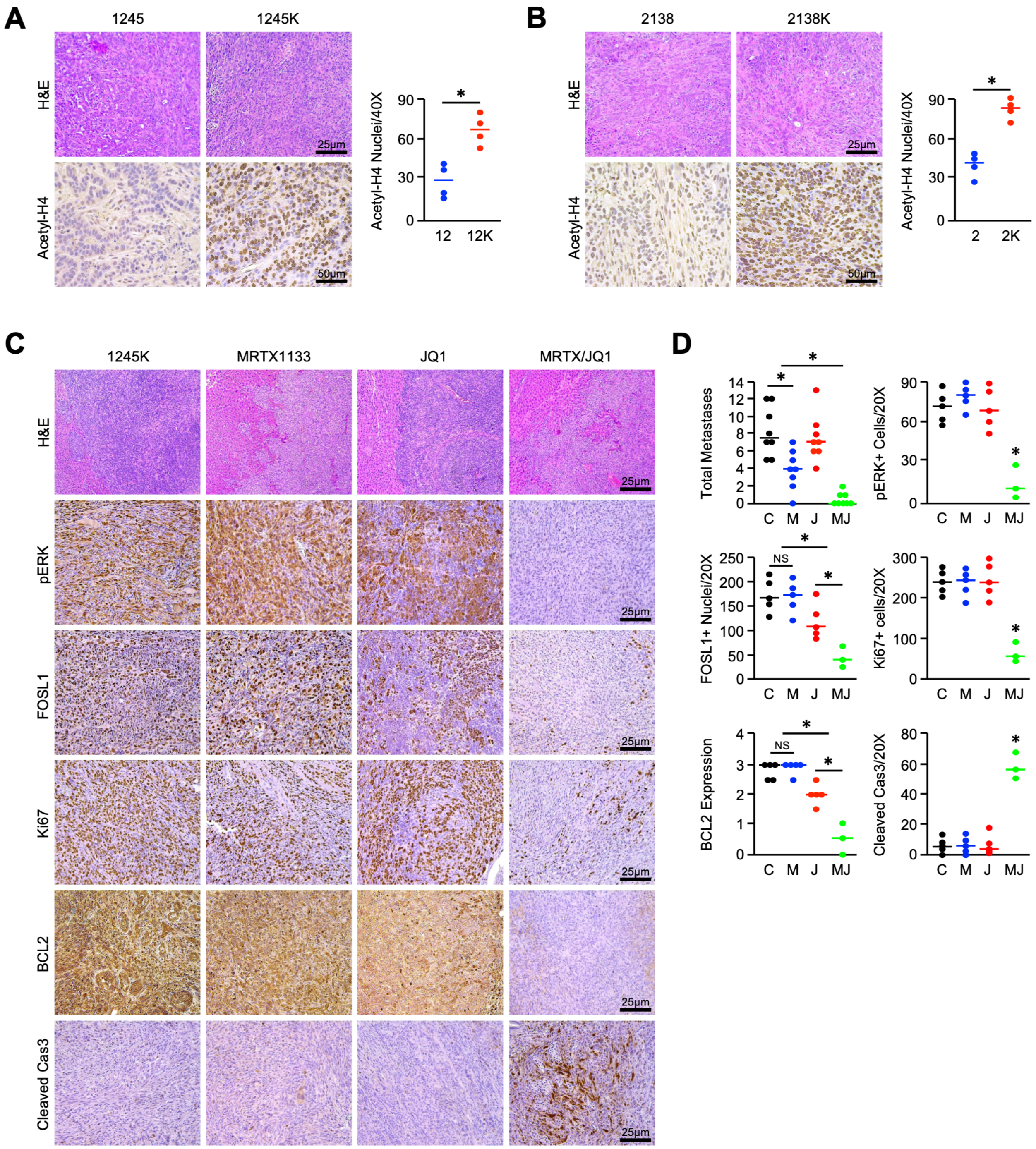


**Figure S7. The combination of MRTX1133 and JQ1 reduces FOSL1 and BCL2 expression in metastatic lesions in the 1245K tumor model**

**(A,B)** 1245, 1245K, 2138, or 2138K cells were collected, resuspended in a 1:1 mixture of Matrigel and full serum media, and 25,000 cells were injected into the flanks of background matched C57BL/6 mice. Mice were euthanized when tumor volume exceeded 1200-1500 mm$^3$, ulcerated, or when mice showed clear signs of health decline, e.g., weight loss, ascites, or lethargy. At the study endpoint, tissues were collected and stained with H&E or via immunohistochemistry for acetyl-H4, quantified as described, and results displayed as individual value plots. **(C,D)** 1245K cells were collected, resuspended in a 1:1 mixture of Matrigel and full serum media, and 10,000 cells were injected into the pancreas of background matched C57BL/6 mice. Mice were allowed to develop a ~250-300 mm$^3$ tumor, at which point they were enrolled into one of four treatment groups. Mice were treated with daily injections of DMSO (vehicle control), 30 mg/kg MRTX1133, 50 mg/kg JQ1, or a combination of MRTX1133 and JQ1 (N = 8 mice/group). Mice were euthanized either upon ulceration of the skin or when showing clear signs of health decline, e.g., weight loss, ascites, or lethargy. At the study endpoint, metastatic lesions from the liver were sectioned and stained with H&E or via immunohistochemistry for pERK, FOSL1, Ki67, BCL2, or Cleaved Caspase 3, quantified as described, and results displayed as individual value plots. (C: DMSO Control, M: MRTX1133, J: JQ1, MJ: MRTX1133/JQ1, *$p < 0.05$).

### Figure S8

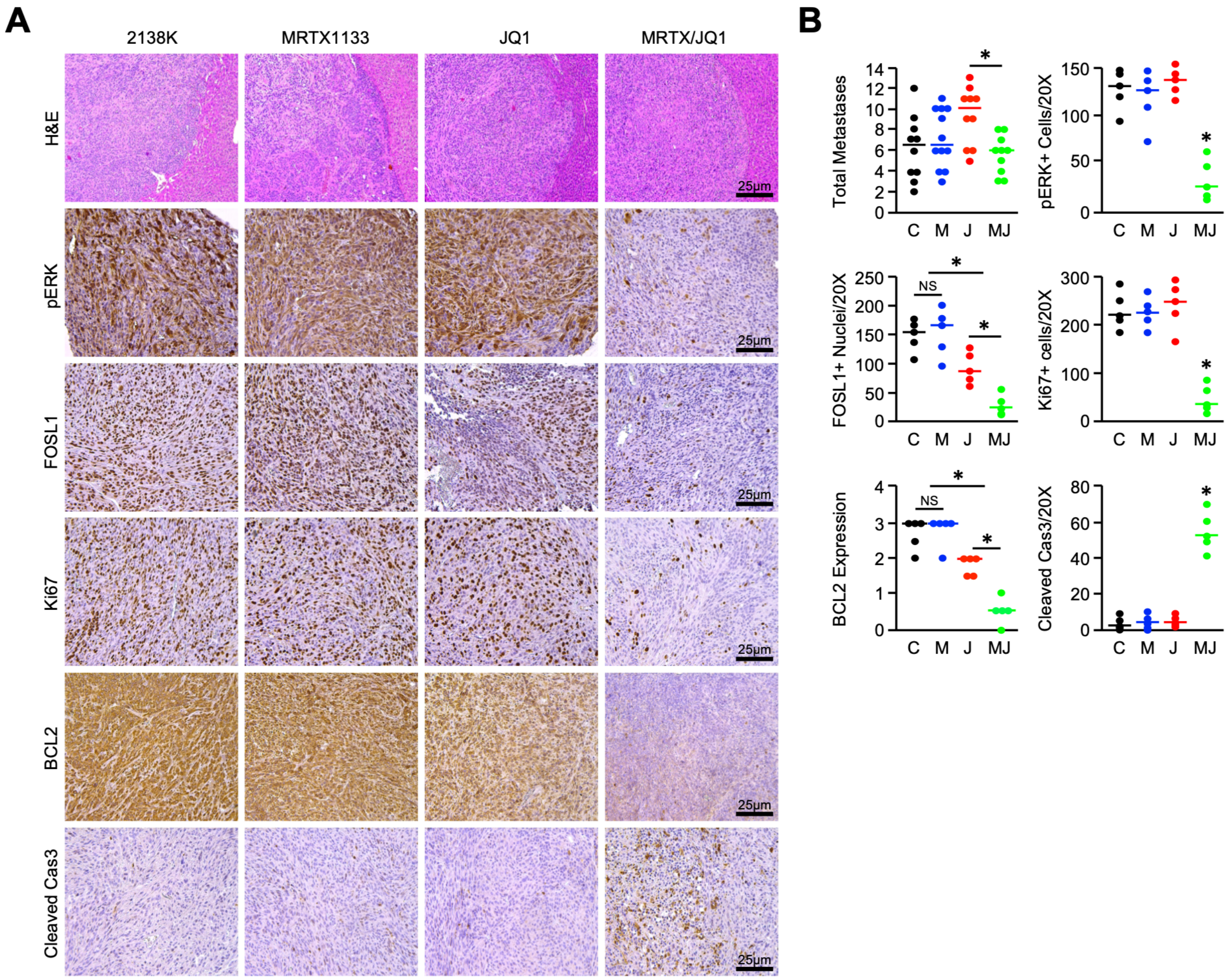


**Figure S8. The combination of MRTX1133 and JQ1 reduces FOSL1 and BCL2 expression in metastatic lesions in the 2138K tumor model**

**(A,B)** 2138K cells were collected, resuspended in a 1:1 mixture of matrigel and full serum media, and 7,500 cells injected were into the pancreas of background matched C57BL/6 mice. Mice were allowed to develop a ~800-1,000 $mm^3$ tumor, at which point they were enrolled into one of four treatment groups. Mice were treated with daily injections of DMSO (vehicle control), 30 mg/kg MRTX1133, 50 mg/kg JQ1, or a combination of MRTX1133 and JQ1 (N = 10 mice/group). Mice were euthanized either upon ulceration of the skin or when showing clear signs of health decline, e.g., weight loss, ascites, or lethargy. At the study endpoint, metastatic lesions from the liver were sectioned and stained with H&E or via immunohistochemistry for pERK, FOSL1, Ki67, BCL2, or Cleaved Caspase 3, quantified as described, and results displayed as individual value plots. (C: DMSO Control, M: MRTX1133, J: JQ1, MJ: MRTX1133/JQ1, *$p < 0.05$).
